# Memory consolidation and representational drift

**DOI:** 10.64898/2026.03.09.710554

**Authors:** Denis Alevi, Felix Lundt, Simone Ciceri, Kristine Heiney, Henning Sprekeler

## Abstract

Memory consolidation is the process by which temporary, malleable memories are transformed into more stable, longer-lasting forms. On a coarse anatomical scale, consolidation redistributes memories in the brain, but it remains poorly understood how these changes manifest themselves on the finer, cellular scale of neuronal engrams and how they relate to the cognitive level. In this study, we developed a phenomenological model of engram dynamics under systems consolidation. The model describes consolidation as a brain-wide phenomenon, where memories deterministically follow a trajectory through a space of patterns distributed among brain regions. It captures a broad range of features of memory consolidation, including selective consolidation, semantization, and power-law forgetting. In the model, consolidation is accompanied by population-level changes in neuronal representations that resemble the widely observed phenomenon of representational drift. When only a subset of neurons is observed, the deterministic dynamics of the model can appear stochastic, and a readout of task features deteriorates over time even when a stable readout exists for the full system. Our model offers a dynamical systems perspective on memory consolidation as a distributed process, moving beyond the classic region-centered view, and provides a functional interpretation of drift as a means of redistributing engrams for improved memory retention.

## INTRODUCTION

Memory is the ability to use past experience to influence the present. It relies on neuronal imprints of memories, likely in the form of long-term synaptic changes (Martin and Morris, 2002) and neuronal activity patterns that are reactivated during memory recall (Josselyn and Tonegawa, 2020). The long-term maintenance of memories over months and years requires an active process—memory consolidation—that transforms recent, malleable memories into a stable format (Squire et al., 2015). Memory consolidation occurs both at the synaptic level in the form of molecular changes that increase synaptic stability (Reymann and Frey, 2007, Fusi et al., 2005, Clopath, 2012) and at the systems level in the form of a redistribution of memories across different areas of the brain (Squire and Alvarez, 1995). Both forms are thought to address the plasticity-stability dilemma, i.e., the challenge of both learning rapidly and retaining memories for long periods of time (McClelland et al., 1995, Fusi et al., 2005, Roxin and Fusi, 2013, Remme et al., 2021).

Systems consolidation redistributes memories across different brain regions, but the accompanying changes at the cellular level are not well understood. The act of remembering appears to rely on “engram cells” (Josselyn and Tonegawa, 2020, Han et al., 2009, Liu et al., 2012) that are consistently reactivated during memory recall over long time spans after memory acquisition. On the other hand, memories that are initially dependent on the hippocampus gradually lose this dependence, suggesting that the engram should change and gradually transform into a brain-wide “engram complex” of linked ensembles of engram cells (Roy et al., 2022), potentially involving the spontaneous reactivation of engrams during sleep (Diekelmann and Born, 2010).

While the dynamics of engrams as the neuronal representation of memories remain unclear (Zaki and Cai, 2024), the dynamics of neuronal representations of space and other task variables have recently been studied extensively. Even after a task has been successfully learned, the associated representations continue to change over the course of days to weeks (e.g., Ziv et al., 2013, Driscoll et al., 2017, Rule et al., 2019, Schoonover et al., 2021). This phenomenon, termed “representational drift”, is experience-dependent (Geva et al., 2023, Khatib et al., 2023) and therefore also relies at least partially on neuronal reactivation. Mechanistic theories of representational drift have suggested that it results from continued learning (Driscoll et al., 2022, Devalle et al., 2025) or stochastic synaptic fluctuations (Qin et al., 2023, Eppler et al., 2026). Yet, whether drift is merely a stochastic epiphenomenon that can be compensated for (Rule and O’Leary, 2022, Micou and O’Leary, 2024) or whether it is a directed, purposeful process is unclear (Micou and O’Leary, 2023).

The goal of the present study is to provide a conceptual bridge between the classical, area-centered view of systems consolidation and the cellular level of neuronal engrams. To this end, we develop a phenomenological model for the dynamics of neuronal engrams during consolidation. The model is mathematically similar to a recurrent neuronal network, allowing us to reinterpret various dynamical phenomena known from neural networks in the domain of memory consolidation. It provides an engram-based view of psychological phenomena, such as selective forgetting and memory semantization, and reproduces power-law forgetting without the need for neuron- or area-specific learning rates (Roxin and Fusi, 2013). Finally, we show that the dynamics of memory consolidation are accompanied by representational changes that share many hallmarks of representational drift, suggesting that representational drift may be a side effect of an active redistribution of memories for long-term maintenance.

## RESULTS

### A population model of memory engram dynamics

We consider a set of memories that can be reactivated by associated cues. Each memory is represented in the brain by an engram vector **e**_*µ*_ (*µ* = 1, 2, …) that describes the participation of different neurons in the *µ*-th memory (Fig. 1A). We do not link this “participation” to a specific physiological signal but rather consider it a conceptual variable that may correlate with a variety of measurements, such as neural activity or immediate early gene expression.

**Figure 1:**
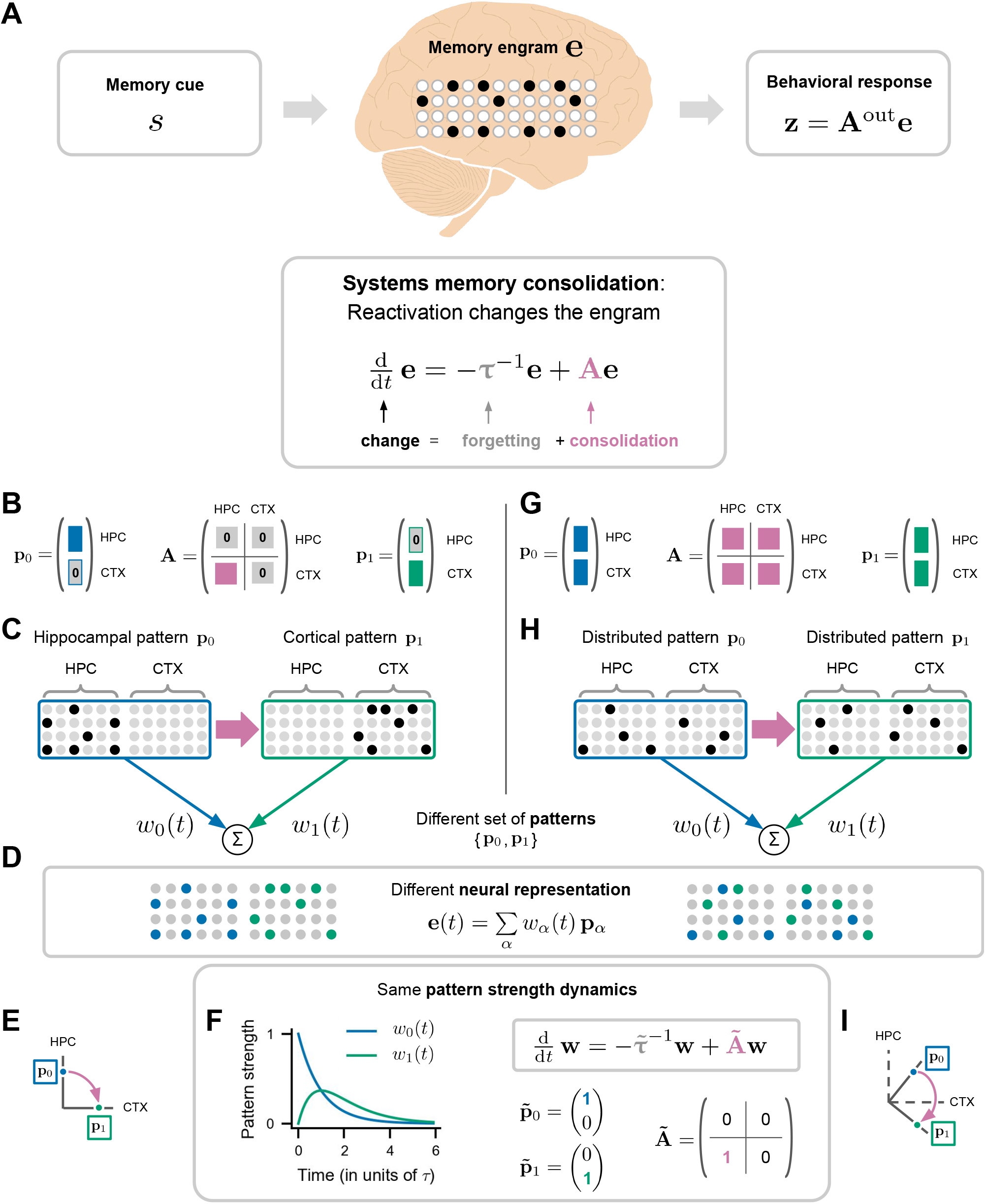
Systems consolidation as sequential dynamics. **A**. Model for the dynamics of a cue-driven memory engram **e. B**. Classical view of memory consolidation: A consolidation matrix **A** with entries that only consolidate from hippocampus (HPC) to cortex (CTX) consolidates a hippocampal engram pattern pattern **p**_0_ into a cortical engram pattern **p**_1_. **C**. Illustration of the consolidation (pink arrow) between the hippocampal (blue) and cortical (green) engram pattern. **D**. The neural engram dynamics **e**(*t*) can be described as a weighted sum of the elemental patterns **p**_*α*_ and time-varying pattern strengths *w*_*α*_(*t*). **E** Illustration of the orthogonality of the hippocampal and cortical patterns. **F** Simulation of the pattern strength dynamics with same pattern forgetting time *τ* for both patterns. **G**,**H**. Same as **B**,**C** but for the consolidation between two distributed (“brain-wide”) engram patterns. **I**. Illustration of the hidden sequential dynamics between distributed, orthogonal patterns.

The model aims to describe how the engrams **e**_*µ*_ change over time. It rests on four simplifying assumptions (for a detailed derivation, see Supplementary Section S1). First, changes in an engram are driven by its reactivation, such that the change in the engram depends on the shape of the engram itself. Second, the reactivation of a given engram does not change other engrams, such that we can describe the dynamics of an engram independently of all the others. Third, we assume that engram changes can be separated into a forgetting component and a component that describes the active consolidation of the memories. Finally, we approximate the engram dynamics as linear. These assumptions lead to the following engram dynamics:

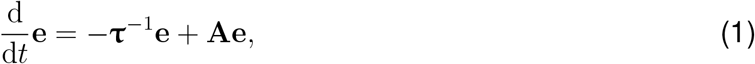

where **τ** is a diagonal matrix of forgetting times for each of the neurons, and **A** is a consolidation matrix that describes how the participation of any neuron in the engram is redistributed among the other neurons during systems consolidation. For the sake of simplifying notation, we start with the dynamics of a single memory and thus exclude the memory subscript *µ*.

We assume that the recall of the engram generates a behavioral readout that we assume to depend linearly on the engram:

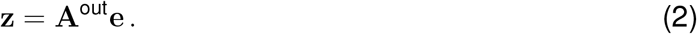

Here, **z** is a readout variable representing, for example, subjective recall or a behavior in response to the elicited memory, and **A**^out^ is the readout matrix. The linear picture is likely a relatively crude approximation of both the dynamics and the readout of the memory engrams, but it provides a geometric perspective on memory consolidation that is conceptually useful.

The computational model in Eqs. (1) and (2) is mathematically equivalent to a linear recurrent neural network (RNN) but with crucial differences in interpretation (Table 1). The primary difference is that our model focuses on the evolution of engram participation over time, rather than the firing rate dynamics typically modeled in RNNs. The consolidation matrix thus reflects the influence neurons have on each others’ involvement in encoding memory, rather than the synaptic connections of the traditional RNN weight matrix. The resultant consolidation dynamics also occur on a slower timescale, of hours to years, rather than the millisecond timescales of neural activity.

**Table 1:** Many well-characterized features of traditional RNNs can be reinterpreted in our engram dynamics frame-work.

| Traditional RNN |  | Engram dynamics |
| --- | --- | --- |
| **Definitions** |  |  |
| Neural activity/firing rate | ⟷ | Engram participation |
| Neural time constant (milliseconds) | ⟷ | Consolidation timescales (hours to years) |
| Synaptic weight matrix | ⟷ | Consolidation matrix |
| Initialization | ⟷ | Memory acquisition |
| Neural dynamics | ⟷ | Consolidation |
| **Model dynamics** |  |  |
| Feedforward networks | ⟷ | Unidirectional pattern-to-pattern consolidation |
| Low-rank networks | ⟷ | Selective consolidation & semantization |
| Attractors | ⟷ | Stable engrams |
| Null-space dynamics | ⟷ | Stable memories despite dynamic engrams |

Modeling the dynamics of memory engrams in this way allows us to reinterpret established dynamical phenomena in RNNs in the context of systems memory consolidation. The following sections address in detail the consequences of this reinterpretation of neural network activity as engram dynamics (Table 1).

### Systems consolidation as sequential dynamics

The model allows us to generalize the classic inter-areal view of consolidation to brain-wide representations. First, we present a consolidation matrix **A** that represents the classic view, where initially hippocampal memories transform into cortical ones. We then show how this view can be transformed to a more distributed view, with memories following trajectories in a space of brain-wide engram patterns.

### Classical view: Hippocampus-to-cortex consolidation

In classical models of systems memory consolidation, memories are formed in the hippocampus (HPC) and gradually transferred into cortex (CTX) for long-term storage (McClelland et al., 1995, Roxin and Fusi, 2013). This picture can be encapsulated in our model by splitting the engram into a hippocampal component **e**^HPC^ and a cortical component **e**^CTX^ and limiting the entries of the consolidation matrix **A** to those that transfer memories from hippocampus to cortex (Fig. 1B):

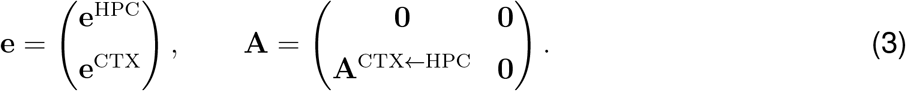

Let us denote the initial memory engram as engram pattern **p**_0_. In the classical view, this initial memory would be exclusively hippocampal (Fig. 1B,C):

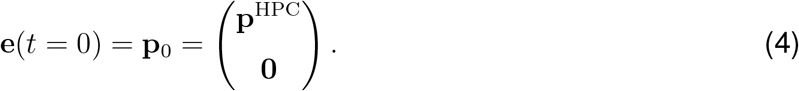

Over time, this initial hippocampal engram pattern **p**_0_ decays as a result of passive forgetting while the consolidation process transforms it into a cortical pattern **p**_1_:

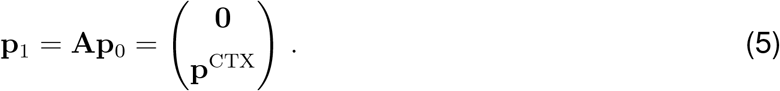

This pattern decays in turn with a time constant that may be slower than hippocampal forgetting (McClelland et al., 1995). Because the hippocampal and cortical patterns are represented in disjoint sets of neurons (Fig. 1D, left), the two patterns are orthogonal (Fig. 1E), and the dynamics take the form of a feedforward network in which each layer transiently stores memories before passing them on to the next layer (Fig. 1F, left).

### Distributed view: Pattern-to-pattern consolidation

To allow a more distributed view of consolidation (Roy et al., 2022), we can move away from the area-centric view of engram dynamics described above, in which a memory transforms explicitly from one part of the brain to another, to a pattern-centric view (Goldman, 2009, Ganguli et al., 2008). In this view, we can retain the previous feedforward dynamics but recast them as transforming an engram within pattern space, where one distributed pattern transforms sequentially into the next.

To introduce the pattern perspective, we represent the time-varying neural engram **e**(*t*) as a superposition of *P* stable patterns of neural engram participation **p**_*α*_, each contributing with a time-varying strength *w*_*α*_(*t*) (Fig. 1D):

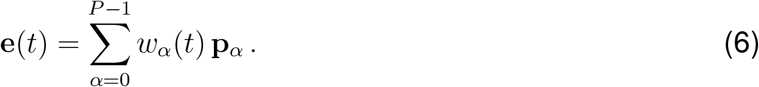

The patterns define the sequential structure of the engram dynamics, while the pattern strengths determine how active each pattern is over time. In this way, each pattern in neural space corresponds to a coordinate axis in pattern space (cf. Fig. 1E,F), and the engram dynamics is fully captured by the pattern strength dynamics:

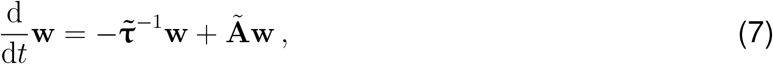

with pattern strength vector **w**(*t*) = (*w*_0_(*t*), *w*_1_(*t*), …, *w*_*P* −1_(*t*)), pattern-specific time constants 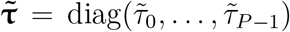 and a pattern-specific consolidation matrix **Ã**(Fig. 1F). We can readily transform between the neural and pattern spaces through an appropriate transform of the forgetting times and consolidation matrix (cf. Supplementary Section S2).

With this pattern-centric perspective, we can generalize the sequential picture of classical interareal consolidation to a sequential consolidation among distributed patterns. To illustrate the idea, consider two patterns **p**_0_ and **p**_1_ that are both distributed across both hippocampus and cortex (Fig. 1G,H). The patterns define the coordinate axes in pattern space 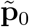 and 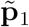, independent of their neural representation (Fig. 1F). A sequential transformation from 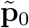 to 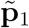 is described by a pattern-specific consolidation matrix with entries limited to those that link these two patterns:

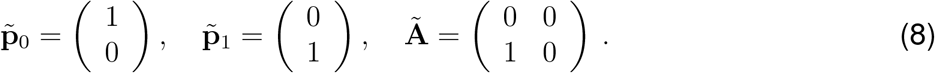

Because we have removed the constraint of the initial pattern being confined to hippocampus, any of the entries in the associated consolidation matrix **A** in the neuron-centric view may be non-zero, effectively mediating a bidirectional consolidation between hippocampal and cortical components of the engram (Fig. 1G).

The two distributed patterns can be seen as rotated versions of the originally hippocampus- and cortex-specific patterns in neural space (Fig. 1I). This rotation preserves the geometry of the engram dynamics and results in a hidden feedforward network, in which each distributed pattern activates the next (Fig. 1F), while individual neurons can participate in multiple patterns. Our model thus transfers the classical picture of hippocampal-to-cortical memory consolidation to the consolidation of potentially brain-wide distributed engram patterns while maintaining the idea of sequential dynamics.

### Selective consolidation through low-rank dynamics

Some memories last a lifetime while others fade quickly, and even those memories that stand the test of time often decline in vividness and detail. Classical models of memory consolidation provide little insight into the neural correlates or mechanisms that underlie such selective forgetting.

In the model, this selective consolidation can be understood as a consequence of the structure of the consolidation matrix. A version of the model that consolidates certain memories while forgetting others can be obtained with a consolidation matrix that is of low rank:

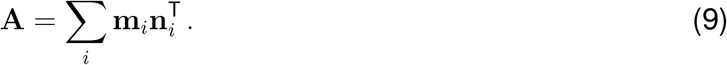

We exemplify this in our hippocampus-to-cortex model (Figs. 2A). This formulation breaks the initial engram pattern **p**_0_ into two components: a selectively retained component that aligns with the selection vectors **n**_*i*_ and a forgotten component that is orthogonal to them (Fig. 2B). The selected component is projected into the consolidated pattern **p**_1_, which is a superposition of the consolidation vectors **m**_*i*_. Thus, only memories that lie in the selection space are consolidated, while memories outside this space are passively forgotten (Fig. 2C).

**Figure 2:**
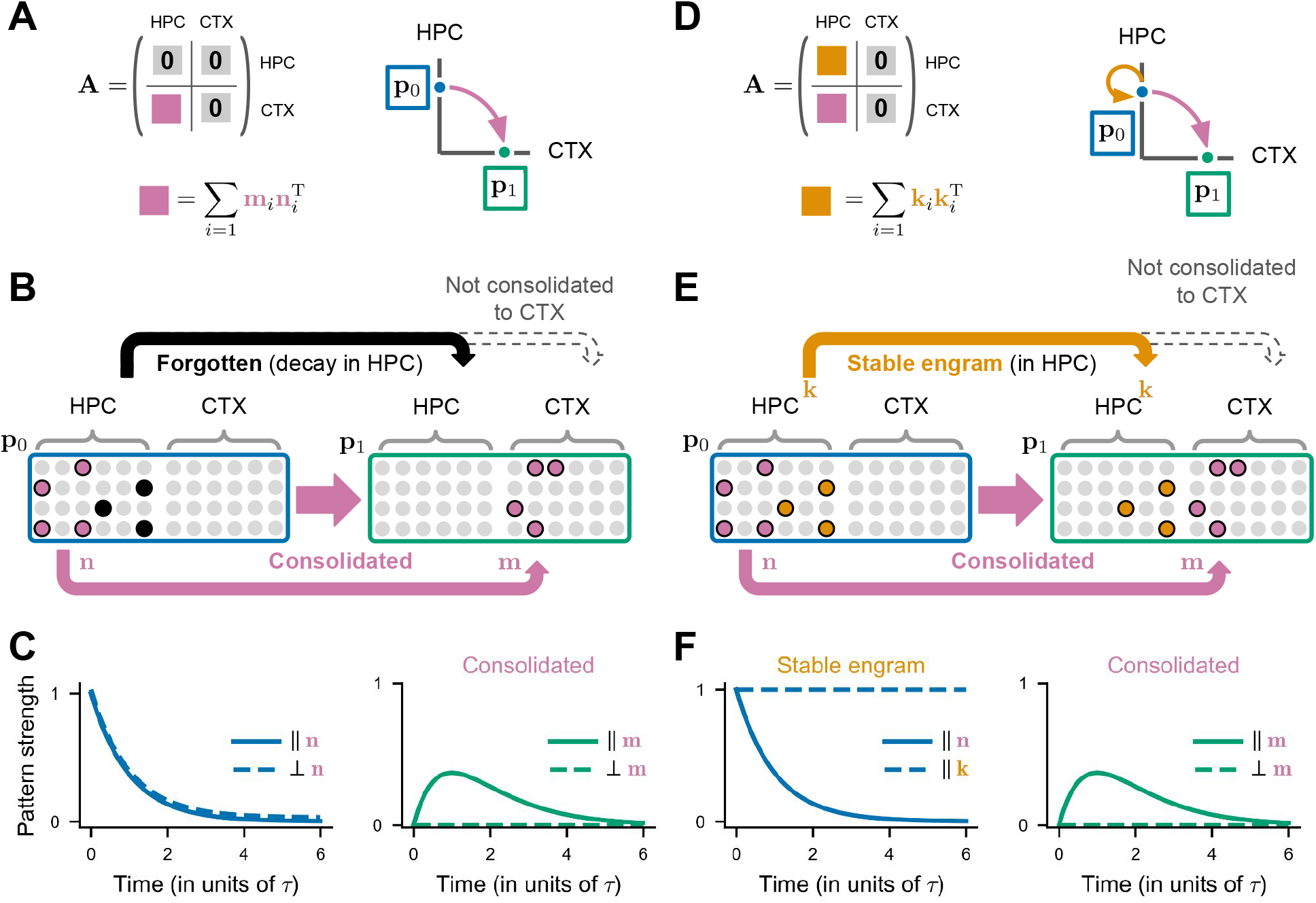
Selective consolidation and stability through low-rank dynamics. **A**. Selective consolidation via a low-rank hippocampus-to-cortex consolidation matrix. **B**. Illustration of low-rank consolidation: One hippocampal component (pink) of the initial engram pattern **p**_0_ (blue) is parallel to the selection vector **n** and is consolidated into a cortical engram pattern **p**_1_ (green) that is parallel to the projection vector **m**. The other hippocampal component (black) is passively forgotten. **C**. Pattern strengths over time with identical forgetting times *τ* . Both hippocampal components are passively forgotten (left, blue lines, small offset for visualization). The engram is consolidated into a cortical engram with a component only in direction of **m** (green, solid). **D-F**. Same as **A-C** but with an additional hippocampus-to-hippocampus consolidation component with identical projection and selection vectors **k** ⊥ **n**, turning the previously non-consolidated hippocampal component (**A**, black) into a stable hippocampal engram (orange).

The selection vectors can select not only which *memories*, but also which *components* of a single memory, are consolidated. If the semantic component of a hippocampal memory aligns with the selection space and the episodic detail is orthogonal, the semantic memory is consolidated into cortex while episodic detail is forgotten.

### Memory stabilization through auto-associative dynamics

Some episodic memories last a lifetime and remain hippocampus-dependent throughout (Winocur and Moscovitch, 2011). This retained hippocampus-dependence can be understood as an active consolidation *within* the hippocampus.

A simple model variant for this phenomenon is obtained when hippocampal selection vectors are paired with themselves as consolidation vectors (Fig. 2D, orange):

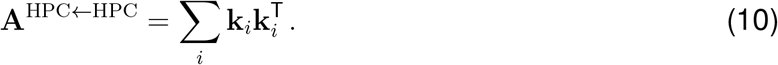

This describes the consolidation of a component of the hippocampal engram into itself over time (Fig. 2E, orange), akin to auto-associative dynamics in RNNs (Hopfield, 1982). When the rate of this “self-consolidation” is balanced with passive forgetting, this results in a stable memory engram in the hippocampus (Fig. 2F, orange).

A stable hippocampal engram component does not preclude a parallel consolidation into cortex. For example, a second selection vector **n** could consolidate a different component of the memory into a cortical pattern **m** (Fig. 2E, F, pink). Thus, an engram can be separated into two parts: one aligned with the stable space **k** that remains in hippocampus, and one aligned with the selection space **n** that consolidates into cortex. Note that it is also possible to maintain hippocampal dependence while also having a dynamic engram within hippocampus; in this case, the selection and consolidation spaces would both be in hippocampus but would not be identical.

### Stable memories despite unstable representations

While the neural representation of a memory changes with consolidation, the memory itself often remains surprisingly stable. The model offers a geometric interpretation for this phenomenon. The number of neurons in the brain is likely very large relative to the dimensionality of memory semantics, i.e., the number of elemental items we can bind together in a memory. This implies that the dimension of the memory readout **z** = **A**^out^**e** (Fig. 1A) is much smaller than that of the engram **e**. As a consequence, many different engram patterns can produce the same readout. The space in which we can change the engram pattern without affecting the readout is referred to as the “output-null space” in the context of RNNs (Kaufman et al., 2014). For the readout to remain stable despite a changing engram, the consolidation dynamics must occur in this output-null space:

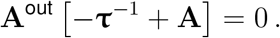

Active consolidation and maintenance are mediated by the consolidation matrix **A**, and its effects can be separated into two components, one in the output-potent and one in the output-null space:

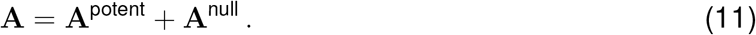

The output-null component is irrelevant for the stability of the memory, because it is by definition unable to change the memory response. Passive forgetting in the output-null space is also inconsequential. The *output-potent* component **A**^potent^ that affects the readout can have two effects. The beneficial effect is that it can counteract passive forgetting and thereby prolong memory lifetimes. However, it can also change the memory response. Take a memory consisting of an episodic and a semantic component that is stored in the hippocampus and decays over time (Fig. 2D). If the hippocampal decay is in the potent space of the memory readout while the dynamics of the semantic component remain in the null-space, the memory will transform from a memory with episodic and semantic components into a purely semantic memory.

### Power-law forgetting of distributed engrams

Theoretical work suggests that a central advantage of memory consolidation is the ability to maintain memories for long periods of time (McClelland et al., 1995, Fusi, 2017). Previous models have achieved slow forgetting, with memory decay following a power law, by sequentially consolidating memories from areas or sets of synapses with short memory lifetimes to areas with increasingly long memory lifetimes (Roxin and Fusi, 2013, Remme et al., 2021, Wixted, 2004).

With our model, we can attain similar gradual power-law forgetting in a system with largely *homogeneous* neural forgetting times, by harnessing the distributed pattern-centric view (Fig. 1F). We endow the patterns **p**_*α*_ that make up the engram **e** with a broad range of forgetting times 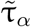. The bulk of patterns rapidly decay, while a few persist for a long time (Fig. 3A). Each pattern consolidates to the next in sequence, then gradually decays (blue, green, grey lines, Fig. 3B). With geometrically distributed forgetting times, these patterns together sum to an engram that decays as a power law (black line, Fig. 3B), by the same mechanism as in earlier models (Fusi et al., 2005, Remme et al., 2021).

**Figure 3:**
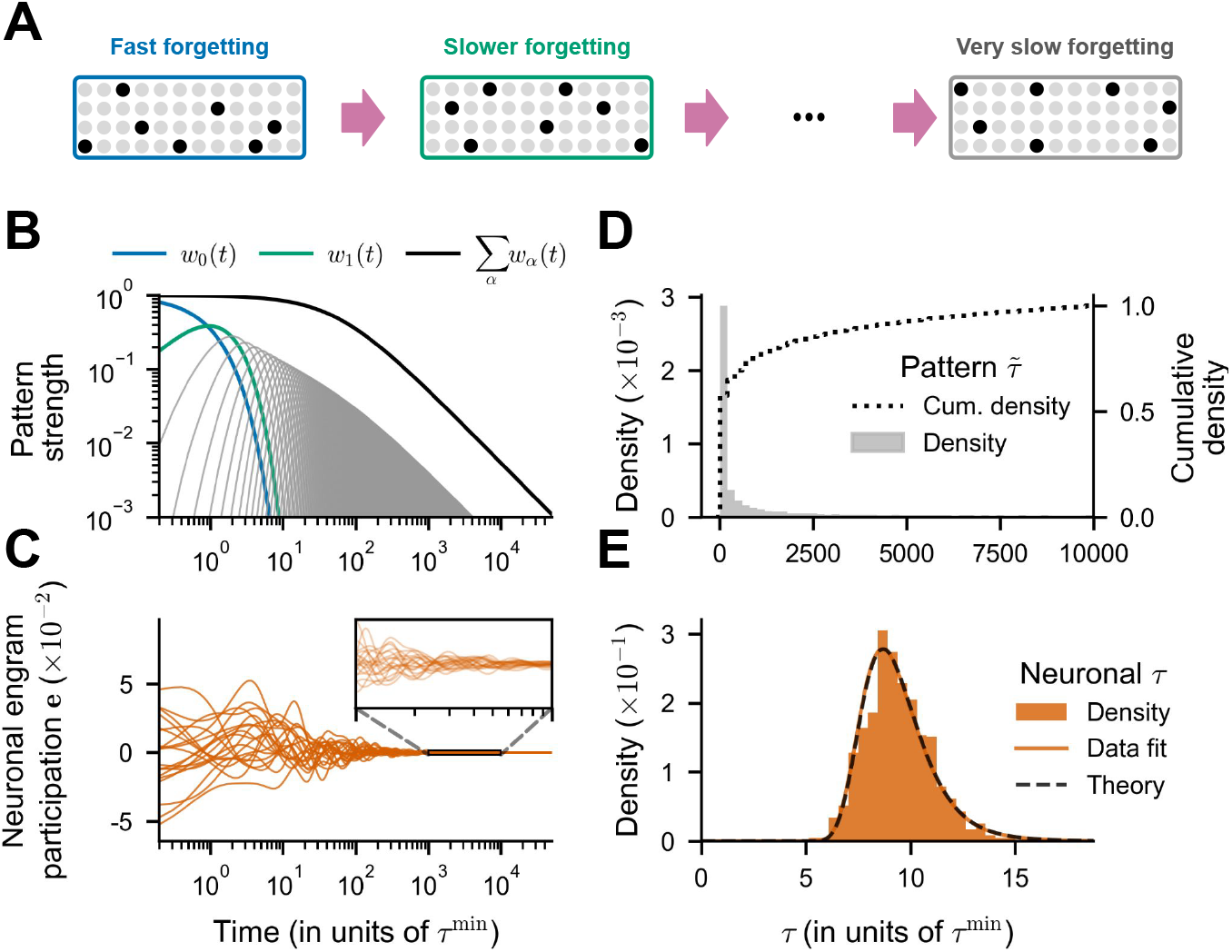
Power-law forgetting from homogeneous neuronal forgetting times. **A**. Schematic of sequential consolidation with distributed pattern forgetting times. **B**. Pattern strengths for all patterns (blue, green, gray lines) with geometrically distributed forgetting times. The readout (black line, sum of all pattern strengths) displays power-law decay. **C**. Neural engram participation for a random subset of neurons, inset showing a zoomed-in version of the same data. **D**. Histogram of pattern forgetting times (gray) and their cumulative density (dotted line). **E**. Histogram of neuronal forgetting times (orange). Lines show the probability density function of a reciprocal normal distribution fitted to the extracted neuronal forgetting times (orange) and computed analytically (black dashed).

Despite these patterns operating on time scales spanning several orders of magnitude, the neural population producing these patterns exhibit more homogeneous forgetting times (CV = 0.164). Because the patterns are distributed, every neuron contributes to a large number of patterns, such that—at least statistically—the engram participation of all neurons decays in the same way (Fig. 3C). While the pattern forgetting times are geometrically distributed (Fig. 3D), the effective neuronal forgetting times follow a reciprocal normal distribution (Fig. 3D,E; Supplementary section S3). This demonstrates that it is possible for a system to achieve power-law forgetting without area- or neuron-specific learning rates.

### Consolidation reproduces key hallmarks of representational drift

While the long-term dynamics of neural representations of memory are not yet well understood, there is a growing body of work on how representations of task variables change over time (e.g., Ziv et al., 2013, Driscoll et al., 2017, Schoonover et al., 2021). This “representational drift” is characterized by gradual ongoing changes in the relationship between neural activity and task variables, such as position or head direction (Rule et al., 2020), without any accompanying change in task performance. Whether this phenomenon is a directed process or the result of stochastic changes remains unclear.

Our model of engram dynamics produces representational changes reflective of those seen in representational drift. To illustrate this, we consider a set of task variables as the cues **s**_*µ*_ in our model. These task variables may be, for example, different visual stimuli or animal locations in the environment. Each cue evokes the associated engram **e**_*µ*_, that is, a representation as a neuronal population vector (Fig. 4A). In the model, these engrams are all consolidated independently with the same consolidation matrix, each following a sequential trajectory through pattern space (Fig. 4B). We again used geometrically distributed forgetting times, producing a slowly decaying memory.

**Figure 4:**
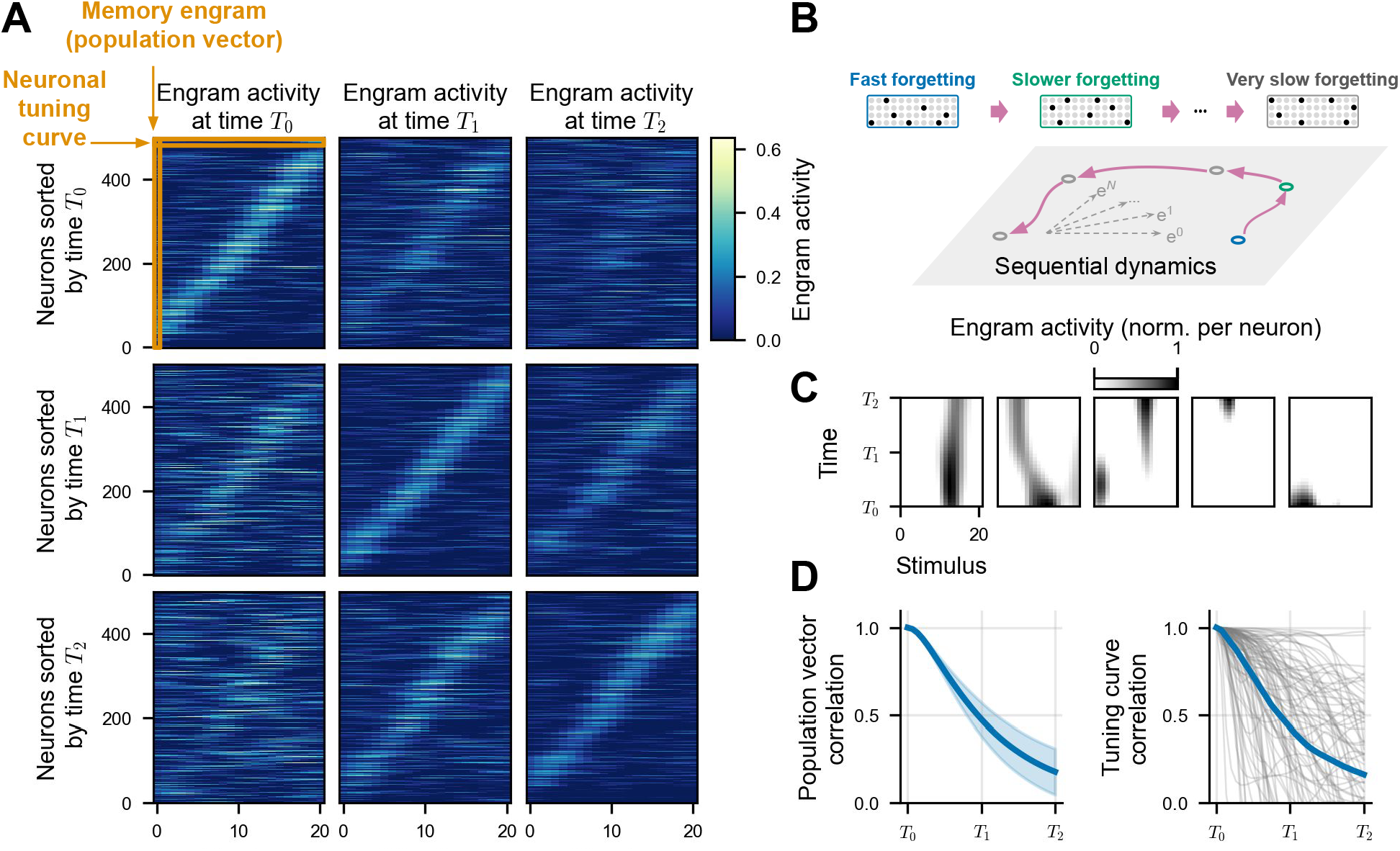
Hallmarks of representational drift. **A**. Stimulus tuning of *S* = 21 engrams at three different time points *T*_0_, *T*_1_, *T*_2_ (grid columns). Each grid row shows same data with neurons sorted by preferred stimulus at a different time point. **B**. Illustration of model of sequential engram dynamics with geometrically distributed pattern forgetting times, same as Fig. 3. **C**. Changes in single neuron tuning for selected neurons, showing stability, gradual drift, abrupt remapping, gain and loss of stimulus-tuning. **D**. Left: Mean population vector correlation of all stimuli. Right: Tuning curve correlations for individual neurons (gray) and their mean (blue).

Over the course of the consolidation, the responses of the neurons to the set of stimuli change, producing changes in their stimulus tuning (Fig. 4C). This is accompanied by a gradual change in the population response, one of the main hallmarks of representational drift (Fig. 4A). When neurons are sorted according to their preferred stimulus at an initial time *T*_0_, their responses tile the stimulus space. At later times, this ordered representation fades, resulting in a gradual decorrelation of the population vectors at each stimulus (Fig. 4D, left). Neuronal tuning curves also decorrelate from their original tuning, albeit at variable rates (Fig. 4D, right), consistent with neural data (e.g., Rule et al., 2020, Geva et al., 2023, Schoonover et al., 2021). Despite these changes, a resorting of the neurons by their preferred stimulus at any time point reveals that the neural responses continue to tile the stimulus space, though with different neurons playing different roles (Fig. 4A). Hence, a deterministic model of engram dynamics can produce key hallmarks of representational drift.

### Representational geometry in subsampled populations

Earlier theoretical work treated representational drift as driven by stochastic fluctuations (e.g., Qin et al., 2023, Delamare et al., 2024). In our model, it results from a deterministic consolidation process. We will now show that this qualitative difference is, unfortunately, difficult to discern in neural data due to subsampling effects: We can usually observe only a fraction of the relevant neurons simultaneously, and engram components in the—typically very large—unobserved neural population gradually consolidate into the observed population (Fig. 5A), where they cause unpredictable changes that cannot be distinguished from stochastic fluctuations.

**Figure 5:**
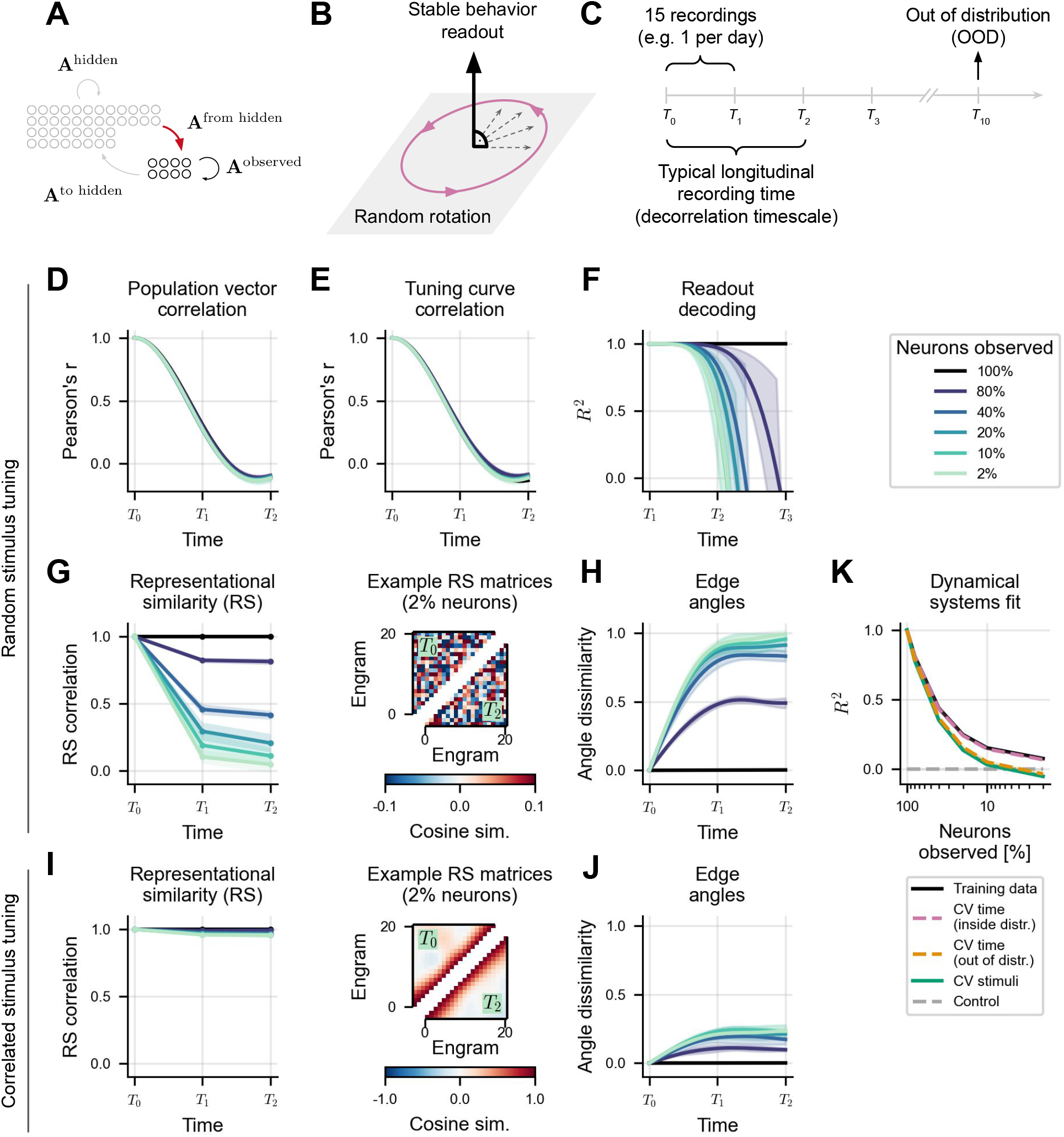
Effects of subsampling. **A**. Subsampling: The hidden population has an unpredictable effect (red arrow) on the dynamics of the observed population. **B**. Model: Engram dynamics in full population perform random rotation in the null-space of a stable behavior readout. **C**. Simulation timescales. Simulation time was chosen to match population vector decorrelation time (see **D**,**E**) and split into 4 time points *T*_0_, *T*_1_, *T*_2_, *T*_3_ with 15 recordings between time points. A later time point was used to evaluate dynamical system fits (see **K**) on out of distribution (OOD) data. **D**. Population vector correlations over time for different fractions of observed neurons. **E**. Tuning curve correlations over time for different fractions of observed neurons. **F**. Linear decoders trained to predict the behavioral readout (see **B**) based on 16 recordings from *T*_0_ to *T*_1_, and evaluated at each recording from *T*_1_ to *T*_3_. **G**. Dynamics of representational similarity over time for random stimulus tuning. Left: Pearson correlation between vectorized upper triangles of cosine-similarity matrices over time for different fractions of observed neurons. Right: Example similarity matrices for 2% neurons observed at time *T*_0_ (upper matrix triangle) and time *T*_2_ (lower matrix triangle). **H**. Edge angle dissimilarity across recordings. **D-H** show results for random stimulus tuning. **I**,**J** Same as **G**,**H**, but for correlated stimulus tuning . **K**. Goodness of fit for linear dynamical systems fitted to simulated data, evaluated on training data within [*T*_0_, *T*_3_] (black line), cross-validated (CV) on left out transitions from the training data (inside distribution, pink dashed line), on a future transition (out of distribution, see **C**, orange dashed line), on left out stimuli (solid green line) and evaluated on a model predicting no change (control, gray dashed line). The number of observed neurons in **D-K** was fixed at *N* = 100, while varying the size of the full population. Shaded areas in all panels show standard deviations across 10 random subsamples.

To study the effects of this apparent stochasticity, we used a variant of the model in which both the behavioral readout and the representational geometry are stable in the full population. Specifically, we confined the engram dynamics to the null-space of a predefined but arbitrary readout variable (Fig. 5B) and restricted the dynamics to high-dimensional random rotations. We then systematically varied the number of unobserved neurons while keeping the size of the observed population and the speed of consolidation fixed (cf. Methods). We initialized engrams with random stimulus tuning and recorded engram activities at evenly spaced time points that cover the decorrelation timescale of the neuronal representation (Fig. 5C). We found that the decorrelation timescales of both the population vector and the tuning curves are independent of the fraction of observed neurons (Fig. 5D,E; Supplementary Fig. S4.1), in line with an analytial derivation using Green’s functions (Supplementary Section S5).

In contrast, subsampling degrades a readout of task variables. To show this, we fit a linear decoder to read out the task variable from the observed component of the engrams during the first half of the simulation and then used this decoder to predict the task variable from the observed engrams over the second half of the simulation. Under full observation of the population, the decoder performance did not degrade over time because observing the full population guarantees access to the stable readout dimension. However, under subsampling, the decoder performance declines over time, with more rapid drops for smaller subsampling fractions (Fig. 5F; Supplementary Fig. S4.1). Thus, subsampling obscures stable task representations that may be present in the full population.

To determine the effect of partial observation on the stability of population geometry, we considered the representational similarity among different stimuli (Deitch et al., 2021) and edge angles in representational space (Schoonover et al., 2021, Sylte et al., 2025). We considered neural representation with random stimulus tuning and with correlated tuning as, e.g., for place cells. Representational similarity measures how similar pairs of stimuli are represented in the population, while edge angle similarity measures the extent to which triplets of stimuli maintain their relative arrangement. Because rotational dynamics preserve the representational geometry, both quantities remain stable during consolidation when the full population is observed (Fig. 5G-J, black lines). The stability of representational similarity under partial observation depends on stimulus tuning. For random tuning, representational similarity becomes increasingly labile with lower sampling fractions (Fig. 5G, left; cf. Schoonover et al. (2021)), while it remains relatively stable for correlated stimulus tuning (Fig. 5I, left; cf. Deitch et al. (2021)). This is because for random stimulus tuning, the representational similarity is determined by random correlations between the representations (Fig. 5G, right), which change under partial observation. However, for correlated stimulus tuning, the representational similarity is predominantly determined by stimulus correlation (Fig. 5I, right), which is much less affected by partial observation. The stimulus tuning also affects the stability of edge angles under partial observation. Independent of tuning, they become increasingly dissimilar with decreasing sampling fraction (Fig 5H,J). However, for correlated stimulus tuning, edge angle dissimilarity saturates at a lower level (Fig. 5H; cf. Sylte et al. (2025)) than for random stimulus tuning (Fig. 5J; cf. Schoonover et al. (2021)).

### Stochastic dynamics in subsampled populations

An obvious test of our model would be to fit it to longitudinal recordings, to uncover the underlying consolidation dynamics. This approach suffers from two challenges. First, such a fit is prone to overfitting. Even for the assumed linear dynamics, the number of data points per neuron (number of stimuli × number of time points) should be at least as high as the number of free parameters per neuron. In the linear consolidation model, there are as many parameters per neuron as there are neurons. This is hard to achieve in longitudinal recordings. Second, even with sufficient data, it will be difficult to predict the influence of the unobserved population on the neural representation. To illustrate this, we fitted linear dynamical systems to simulated data, ensuring sufficient data to constrain the fit. We then evaluated the fitted models on held out stimuli, held out time points within the training data (inside distribution, ID) and observations from a future time point (out-of-distribution (OOD); Fig. 5C). As expected, the fits succeed in the full population and generalize well (Fig. 5K). However, even with relatively minor subsampling, the performance of the fit declined steeply, even on the training data. Hence, the influence of the unobserved population introduces an unpredictable component, such that the dynamics appear stochastic despite their deterministic origin.

## DISCUSSION

In this work, we modeled memory consolidation as slow, reactivation-driven dynamics that reorganizes brain-wide engram patterns over slow timescales of hours to years. By generalizing the classical area-centered view of systems consolidation to a global population-level description, we revealed how established recurrent network motifs can be reinterpreted as mechanisms of long-term memory reorganization. Representational drift emerges as a natural corollary of this global memory reorganization. Finally, subsampling—the observation of a small fraction of the full population—can obscure directed consolidation dynamics, such that they appear random, complicating the recovery of the system dynamics from neuronal recordings.

The suggested dynamical systems perspective on memory consolidation creates a bridge from classical theories of consolidation, which primarily describe the time-course of hippocampal dependency and differ in their postulates for episodic and semantic memories (Squire and Alvarez, 1995, Nadel and Moscovitch, 1997, Winocur et al., 2010), to contemporary views of global memory organization (Moscovitch and Gilboa, 2022). The area-centered view is captured as a special case of a general brain-wide description of memory reorganization. A sustained or fading hippocampal dependency is described by the geometric alignment of memories with the consolidation process, represented by the consolidation matrix *A*: Depending on how an engram aligns with this matrix, it is passively forgotten, actively maintained within the hippocampus, or consolidated into the neocortex. The modeling framework does not provide a natural distinction between episodic and semantic memory and therefore cannot speak to their differential consolidation. However, it can readily describe a selective and partial consolidation such that memories could be stripped of their episodic vividness, rendering them increasingly semantic in nature.

### Mechanistic interpretation & assumptions

The suggested model abstracts away mechanistic details at the cellular and circuit levels and instead focuses on the phenomenological evolution of memory traces. It distinguishes between active consolidation and passive decay, treating both as deterministic drivers of engram evolution. The decay term represents the aggregate effect of stochastic, destructive processes, such as interference from newly stored memories (Robertson, 2012), spontaneous synaptic turnover (Ziv and Brenner, 2018), and neurogenesis (Golbabaei et al., 2025). We note that this formulation captures physical trace erosion rather than forgetting due to transient retrieval failure (de Snoo and Frankland, 2025).

The active consolidation component represents the systematic drive from processes such as circuit-level engram transfer (McClelland et al., 1995, Remme et al., 2021, Bhasin et al., 2024), synaptic plasticity (Lee et al., 2023), assembly formation (Tomé et al., 2024, Golbabaei et al., 2025), and synaptic consolidation (Fusi et al., 2005, Benna and Fusi, 2016), all of which dictate how a memory trace reorganizes over time. The effect of these mechanisms is phenomenologically summarized in the consolidation matrix **A**, so we assumed fixed, linear engram dynamics. This linearity limits the model, because all memories undergo the same dynamics and different aspects of a memory cannot interact nonlinearly. Biological consolidation, however, is adaptive and nonlinear. Experience can reshape the consolidation dynamics themselves in a process known as schema formation (Tse et al., 2007) or meta-learning (Wang et al., 2018, Zhou and Schapiro, 2025). Similarly, the consolidation dynamics often depend on the consolidation context, such as internal states (Chouhan et al., 2020), or the systematic interaction with other memories (McClelland et al., 1995, Schlichting et al., 2015). Our linear framework thus represents a first-order approximation: it captures how memories evolve in a given consolidation context when the underlying consolidation rules are held constant. Generalization to nonlinear dynamics is beyond the scope of the present work.

The assumption that the consolidation of an engram depends on the engram itself amounts to assuming that consolidation is driven by reactivation, because information about the shape of the engram is assumed to be present at the time of consolidation. We remain agnostic as to the type of reactivation involved. In particular, we do not distinguish between replay during sleep (Diekelmann and Born, 2010), wakeful rest (Wamsley, 2022), and the re-experience of a known learning paradigm (Himmer et al., 2019), even though they might contribute differentially to consolidation. Finally, the model treats reactivation merely as a driver of memory reorganization and ignores a potential re-encoding of memory traces during re-experience (Nadel and Moscovitch, 1997), systems reconsolidation (Debiec et al., 2002), or direct interactions between distinct memory traces beyond passive interference (Schlichting et al., 2015).

### Representational drift as a correlate of consolidation

In our model, the same consolidation dynamics that reorganize memory engrams across the brain over years also produce representational drift in local population codes over days. In this view, representational drift is not merely a passive degradation of neural codes that needs compensation (Rule et al., 2020, Kalle Kossio et al., 2021) but a signature of distributed systems consolidation. The model hence unifies the two phenomena by treating the stimulus-behavior mappings in drift studies as cue-response associations that undergo consolidation.

In our model, systems consolidation is deterministic and directed, implying that the associated drift is also directed. Existing models for drift tend to drive it with a stochastic component, attributed to random synaptic updates (Qin et al., 2023, Ratzon et al., 2024, Morales et al., 2024, Eppler et al., 2026), interference from ongoing learning (Devalle et al., 2025), or fluctuations in intrinsic excitability (Delamare et al., 2024, Haimerl and Machens, 2025). In our framework, on the other hand, these same stochastic mechanisms are captured as a passive decay of the engram and do not drive the drift dynamics. Most existing models include an additional directed component in the drift dynamics, e.g. by constraining the random dynamics to a solution manifold (Ratzon et al., 2024, Natrajan and Fitzgerald, 2024), balancing it with plasticity (Qin et al., 2023, Devalle et al., 2025, Morales et al., 2024, Eppler et al., 2026), or by deriving from it a directed area-to-area consolidation component (Kossio and Memmesheimer, 2025). Our model illustrates that a directed component alone—systems consolidation—is sufficient to generate the complex phenomenology of representational drift.

The apparent randomness of drift observed in experiments can be reconciled with deterministic dynamics by accounting for limited observability. When a high-dimensional, deterministic consolidation process is observed through the lens of a small subset of recorded neurons, the resulting dynamics appear dominated by noise and lose the structure and predictability present in the full population. This suggests that random components of drift reported in experimental recordings (Rule et al., 2020, Schoonover et al., 2021) may be the result of limited observability rather than a fundamental property of the neural code.

The model predicts that drift dynamics are deterministic and hence predictable. Unfortunately, the experimental test of this prediction requires overcoming the subsampling barrier (Wilting and Priesemann, 2018, Levina et al., 2022, Qian et al., 2024) to characterize the full consolidation dynamics. For large neural systems such as mammalian brains, this is challenging because longitudinal recordings only observe a fraction of the full brain over a limited time window. Given such limited data, a linear fit of the representational dynamics is prone to overfitting, and even a failure to fit would be a poor falsification of the model, because it may merely be the result of subsampling. However, we expect that fitting may be possible even in large memory systems if the dimension of the neural subspace covered by the relevant engrams over time is small or comparable to the number of recorded neurons. In this case, the unobserved part of the engram does not contain additional information, and the fitting should succeed.

A test of the model could also be within reach for smaller model systems. For example, in the *Drosophila* mushroom body, a small number of mushroom body output neurons represent odor-reward associations (Owald and Waddell, 2015), and their involvement changes through consolidation (Cervantes-Sandoval et al., 2013). While recording the full population of these neurons simultaneously is not currently possible in Drosophila, this or other model systems may, in the future, offer the required access to study whether memory consolidation, engram dynamics and representational drift are different sides of the same coin.

## RESOURCE AVAILABILITY

### Data and code availability

All code will be made publicly available upon publication.

## ACKNOWLEDGMENTS

This work was supported by the Deutsche Forschungsgemeinschaft (DFG, German Research Foundation) under the Collaborative Research Center SFB 1315 (project number 327654276) and SFB/TRR 384 “IN-CODE”. KH is funded by an Alexander von Humboldt Research Fellowship.

## AUTHOR CONTRIBUTIONS

Conceptualization DA, FL, HS; methodology DA, FL, SC, HS; investigation DA, FL, SC, HS; writing—original draft DA, FL, SC, KH, HS; writing—review & editing DA, SC, KH, HS; funding acquisition KH, HS; supervision KH, HS

## SUPPLEMENTAL INFORMATION INDEX

Supplementary section S1: Model derivation

Supplementary section S2: Transforming between neural space and pattern space

Supplementary section S3: Mathematical derivation of power-law forgetting with homogeneous neuronal dynamics

Supplementary figure S4.1: Effects of subsampling for correlated stimulus tuning

Supplementary section S5: Mathematical derivation of apparent stochasticity under subsam-pling

## METHODS

All simulations used forward Euler integration with time step Δ*t* = 0.001 for Figs. 1,2,4,5 and Δ*t* = 0.1 for Fig. 3. Figs. 1–3 were simulated in pattern space (Eq. 7), where the *i*-th pattern 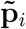 was chosen as the *i*-th standard basis vector in *P* dimensions, i.e. with components 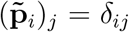, where *δ* is the Kronecker-Delta. Figs. 4 and 5 were simulated in neural space (Eq. 1).

### Systems consolidation as sequential dynamics

For Fig. 1 and 2, forgetting times were the same for all patterns 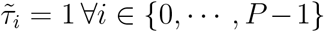. For Fig. 1 we simulated *P* = 2 patterns, initialized the engram with the first pattern 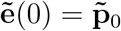 and used consolidation matrix

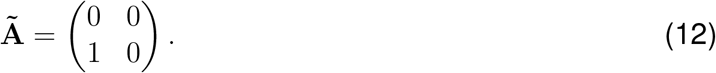

For Fig. 2, we simulated *P* = 4 patterns (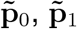 in hippocampus and 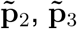 in cortex), initialized the engram with both hippocampal patterns active 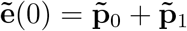. The consolidation matrix was

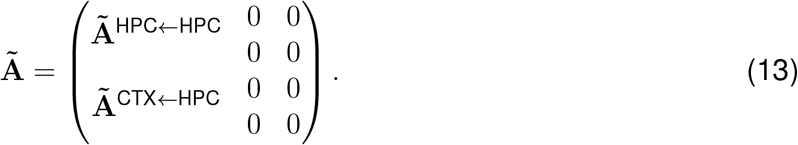

For the selective hippocampus-to-cortex consolidation, the first hippocampal pattern was consolidated via selection vector 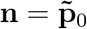 and projected into the first cortical pattern via 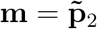, such that

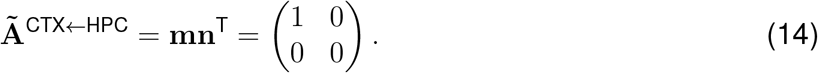

For the decaying second hippocampal component, we set **Ã**^HPC←HPC^ = **0**. For the stable second hippocampal component, we set 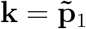, such that

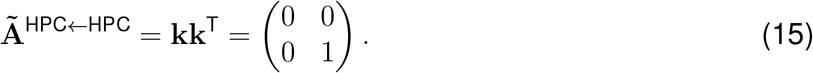

### Power-law forgetting from homogeneous neuronal forgetting times

For Fig. 3, we simulated *P* = 500 patterns with geometrically distributed forgetting times 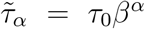 for *α* ∈ {0, …, *P* − 1}, with *τ*_0_ = 1, *β* = (*τ*_max_*/τ*_0_)^1*/*(*P* −1)^, and *τ*_max_ = 10000. Consolidation was implemented as sequential pattern-to-pattern transfer via an off-diagonal pattern consolidation matrix 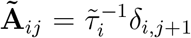. The readout was computed as a sum of all pattern strengths via readout vector **Ã**^out^ = (1, 1, …, 1)^T^. Neural engram pattern entries (**p**_*i*_)_*j*_ for neuron *j* in pattern *i* were sampled from a standard normal distribution in ℝ^*N*^ where *N* = 1000 was the number of neurons. Each pattern **p**_*i*_ was then normalized to have length 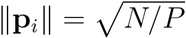, such that the entries in the engram participation vector **e** where independent of the number of patterns *P* and number of neurons *N* . The pattern space dynamics were transformed to neural space via **e**(*t*) = **Pw**(*t*), where **P** ∈ ℝ^*N×P*^ was the matrix of neural engram patterns and **w** the vector of pattern strengths. The effective time constants of individual neurons were extracted from the diagonal entries of the neural consolidation matrix 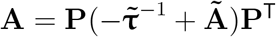, giving *τ*_*i*_ = −1*/***A**_*ii*_ for each neuron *i*.

The theoretical distribution of neural time constants was computed as the reciprocal normal distribution

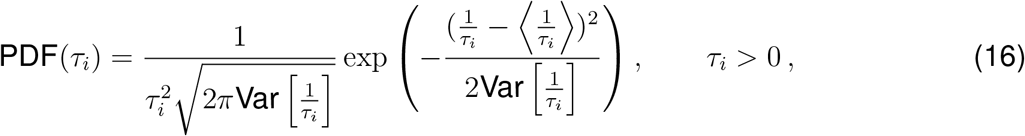

with

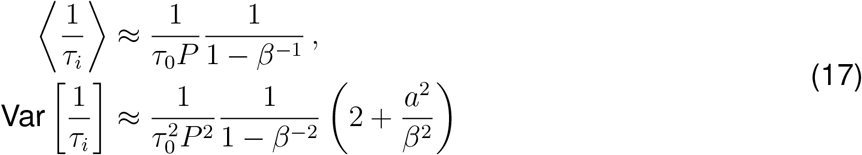

and *a* = 1. For analytical derivations, see Supplementary Section *S*3.

### Representational drift simulations

For Fig. 4 and 5, we simulated engram dynamics for *S* = 21 stimulus-evoked engrams. Simulations were run for a burn-in period *t*_burn_ after which recording time began as *t* = 0 and ran until *t* = *t*_sim_. Population vectors were recorded at *k* evenly spaced time points *t*_0_, *t*_1_, …, *t*_*k*−1_ within the total simulation time. We defined time points *T*_0_, *T*_1_, *T*_2_ as recording indices *t*_0_, *t*_15_, *t*_30_, respectively. For Fig. 5, we defined an additional time point *T*_3_ as recording index *t*_45_. Simulation times were chosen such that population vector correlation dropped to approximately zero from *T*_0_ to *T*_2_ (Fig. 4: *t*_sim_ = 30 with *k* = 31 in [*T*_0_, *T*_2_]; Fig. 5: *t*_sim_ = 3 with *k* = 46 in [*T*_0_, *T*_3_]). Population-vector and tuning-curve correlations were computed as Pearson correlations relative to *t*_0_.

### Hallmarks of representational drift

For Fig. 4, engrams were initialized with mixed Gaussian tuning by tiling the stimulus space with Gaussian tuning curves (*σ* = 2.1) and applying a random orthogonal mixing across neurons. The temporal evolution of each engram was independently simulated using the same sequential pattern-to-pattern dynamics with geometric time-constant distribution as for Fig. 3, but with *N* = 500 neurons, *P* = 500 patterns, and *τ*_max_ = 15. We simulated *t*_burn_ = 10 time units of burn-in before recording population vectors. We processed the simulated activity traces by rectifying negative values to zero. This corresponds to simulating sub-threshold neuronal activity that is only observable when it crosses the spiking threshold (Mainali et al., 2025). For visualization of single neuron tuning curves (Fig. 4C), we normalized each neuron’s activity across stimuli and time to [0, 1]. Activities in Fig. 4A were not normalized.

**Figure 5: Subsampling, decoding, and fits of the dynamics**. Engrams with random stimulus tuning were initialized with normally distributed engram participation entries, sampled from 𝒩 (0, 1). Engrams with correlated stimulus tuning were initialized using the mixed Gaussian tuning used for Fig. 4 (with *σ* = 2.1). Stable rotational dynamics were constructed by sampling a random matrix **G** with entries 𝒩 (0, 1*/N*_neurons_), forming an antisymmetric generator **A**_0_ = **G** − **G**^T^. We then set the first dimension of **A**_0_ to zero and defined the readout matrix 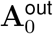 to read out from this dimension, such that the dynamics lie in the nullspace of the readout. Finally, we mixed this dimension across neurons by conjugating both matrices by a random orthogonal matrix **O** to get **A** = **OA**_0_**O**^T^ and 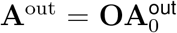. We fixed the number of observed neurons at *N*_obs_ = 100 and varied the sampling fraction *f* ∈ {0.02, 0.1, 0.2, 0.4, 0.8, 1.0} by simulating larger networks with *N*_tot_ = ⌈*N*_obs_*/f*⌉ and randomly selecting *N*_obs_ neurons for analysis. All results were averaged across 10 independent neuronal samples from the same full population. Each simulation consisted of *t*_burn_ = 5 of burn-in time, followed by *t*_sim_ = 3 simulation time sampled at 46 time points. We defined time points *T*_0_, *T*_1_, *T*_2_, *T*_3_ as recording indices *t*_0_, *t*_15_, *t*_30_, *t*_45_, respectively. Decoding used ordinary least squares to predict **z**_*µ*_(*t*_*k*_) from the observed activity **e**_*µ*_(*t*_*k*_), training on all stimuli and the first *k* = 16 time points (in [*T*_0_, *T*_1_], i.e. trained on *S* × *k* = 21 × 16 = 336 samples) and reporting *R*^2^ across time in [*T*_1_, *T*_3_] (normalized such that a model always predicting the average of the data gives *R*^2^ = 0; default from `sklearn.metrics.r2_score`). Representational similarity matrices were computed as cosine similarity between engram vectors across neurons at *T*_0_, *T*_1_, *T*_2_ and compared by correlating their vectorized upper triangles (Deitch et al., 2021). Population geometry was quantified by edge-angle matrices (Schoonover et al., 2021, Sylte et al., 2025) summarized by the Frobenius norm ∥**M**(*t*_*k*_) − **M**(*t*_0_)∥_*F*_, where **M**(*t*) is a *S* × (*S* − 2)(*S* − 1)*/*2 dimensional matrix with all edge angles connecting to a single engram vector per matrix row. Edge angle matrices were normalized by a shuffle baseline obtained by permuting engram labels (edge angle matrix rows) at *T*_2_. To fit dynamics from subsampled data we used ordinary least squares regression without intercept to learn a one-step increment model Δ**e**_*µ*_(*t*_*k*_) = **e**_*µ*_(*t*_*k*+1_) − **e**_*µ*_(*t*_*k*_) from **e**_*µ*_(*t*_*k*_) (training on all consecutive transitions in the training window) and report training *R*^2^, 5-fold cross-validated *R*^2^ across held-out stimuli, leave-one-out cross-validated *R*^2^ across held-out time transitions within the training window, a control baseline predicting Δ**e**_*µ*_ = 0, and an extrapolation score on a future out-of-distribution (OOD) transition ending at *T*_10_. Fits were performed on the random stimulus tuning simulations using 46 recordings, but with 500 simulated engrams to make sure all fits were constrained by sufficient data.

## SUPPLEMENTARY MATERIAL

### S1 Model derivation

The suggested model of memory consolidation describes the change in engrams over time as a linear dynamical system. This section gives a detailed derivation of this model.

In our model, each of a set of *M* cues or stimuli *s*_*µ*_ (*µ* ∈ {1, …, *M*} ) is associated with a corresponding memory engram **e**_*µ*_ describing the participation of a population of *N* neurons. Each engram **e**_*µ*_ is given as an *N* -dimensional vector 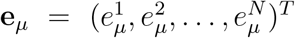 with elements 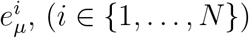 giving the participation of each neuron in the population.

#### Assumption 1: Reactivations drive dynamics

First, we assume that changes in engrams are a function of the engrams themselves. This assumption is based on the findings that the disruption of replay during sleep impairs memory consolidation (Girardeau et al., 2009) and cue reactivation strengthens memories (Rasch et al., 2007), suggesting memory reactivation drives changes in the engram. This can be described by

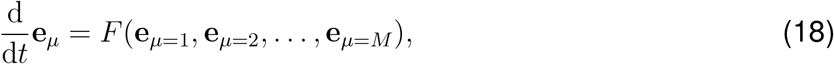

where *F* is a function of all reactivated memory engrams. This formulation represents the case in which engrams may be influenced by the reactivation of all other engrams; that is, reactivation of a given memory can influence engrams of not only that memory but also all other memories.

#### Assumption 2: Non-interference of engrams

We next impose the constraint that the dynamics of a given engram **e**_*µ*_ are influenced only by the reactivation of the engram itself and not by any of the other engrams represented in the system. This assumption decouples the dynamics of the engrams, as

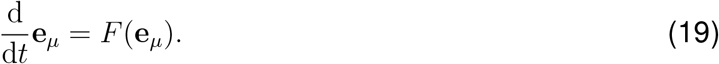

This allows us to consider the dynamics of each engram independent of all other engrams in the system.

#### Assumption 3: Decomposition into forgetting and consolidation terms

Changes in the engram involve both the redistribution of neural participation among the population and gradual decay due to forgetting or interference with newly encoded memories. We here assume that these two components can be treated as separate terms in *F*, as

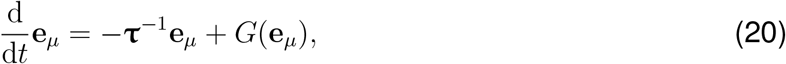

where **τ** = diag(*τ*_1_, *τ*_2_, …, *τ*_*N*_ ) is a diagonal matrix of forgetting times for each of the *N* neurons and *G* is a function describing the redistribution of the engram.

#### Assumption 4: Linearity

As a first approximation, we assume *G* takes a linear form:

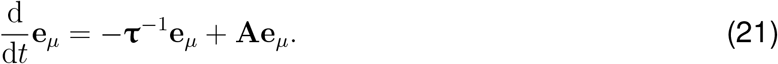

We also assume linearity in reading the memory out from the system:

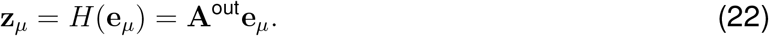

Although this assumption of linear *G* and *H* is quite strong, this first-order approximation provides a geometric perspective on engram dynamics that yields valuable insights into the system.

### S2 Transforming between neural space and pattern space

Our model describes engram dynamics as a sequential reconfiguration of the engram from pattern to pattern. In the classical view of hippocampus-to-cortex consolidation, these patterns **p**_0_ and **p**_1_ are each confined to a single brain region (Eqs. 4 and 5):

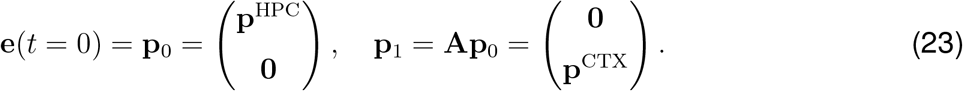

Here, **p**_0_ is the initial engram pattern, located entirely within the hippocampus, and **p**_1_ is the engram pattern that emerges in cortex during consolidation. In this formulation, neurons are tied to a single area-specific pattern, and the neural dynamics are thus explicitly tied to the pattern dynamics.

We release this constraint by viewing consolidation as a distributed process, whereby engrams traverse a trajectory in the space of patterns **p**_*α*_ ∈ ℝ^*N*^ not confined to specific brain regions or populations of neurons. This distributed view also means that a single neuron may participate, to varying degrees, in multiple patterns. The engram is then given by (Eq. 6):

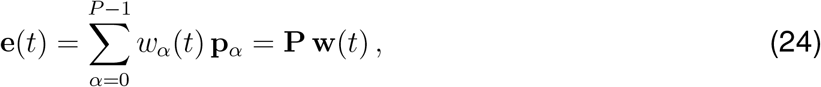

where *w*_*α*_(*t*) are the time-varying pattern strengths linking the engram to the patterns **p**_*α*_ and **P** ∈ ℝ^*N×P*^ is the pattern matrix consisting of all possible patterns.

We can then describe the engram dynamics in this pattern-centric coordinate system by focusing on the dynamics of the pattern strengths **w** (Eq. 7):

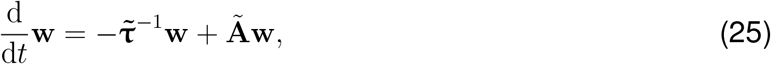

where the time constants 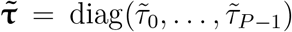 and consolidation matrix **Ã**are described in pattern space rather than neural space.

We can transform these pattern dynamics back to neural space with a suitable transformation of the consolidation matrix. The dynamics in neural space are given by

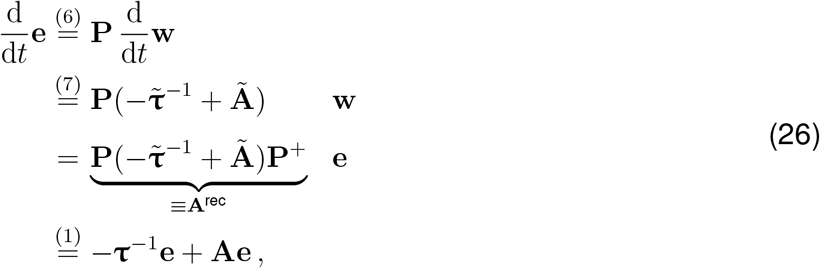

where **P**^+^ is the pseudoinverse of **P**. We interpret the diagonal entries of the matrix **A**^rec^ as inverse time constants **τ**^−1^ and the non-diagonal entries as the consolidation matrix **A** in neural space.

To keep the readout the same when moving between neural and pattern spaces, we must also transform the readout matrix:

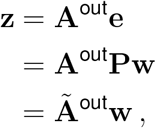

From this, the readout **Ã**^out^ from the pattern strengths and that **A**^out^ from the engram are related as **Ã**^out^ = **A**^out^**P**.

### S3 Mathematical derivation of power-law forgetting with homogeneous neuronal dynamics

Power law forgetting is achieved by a consolidation matrix that represents a feedforward chain of *P* patterns with geometrically distributed pattern forgetting times:

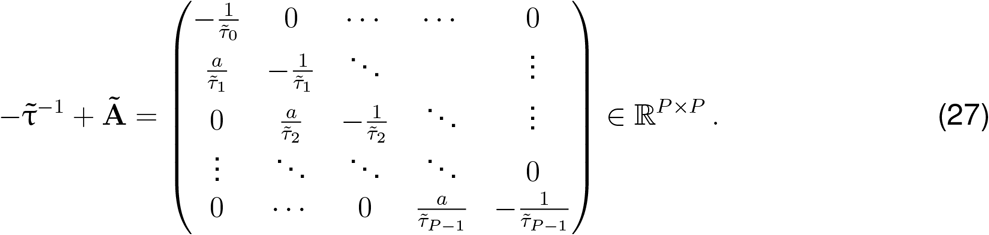

where we scaled the feedforward coupling between the patterns with the inverse forgetting times to avoid uncontrolled growth of the pattern strengths for later patterns. For geometrically distributed pattern forgetting times, we choose

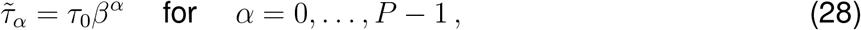

with a time scale *τ*_0_ and a factor *β >* 1 that controls the rate at which the forgetting times increase with the pattern number *α*.

To find the effective timescales of the dynamics of individual neurons, we transform the dynamics in pattern space back into neural space (Eq. 26) and analyze the diagonal entries

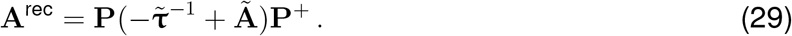

In the limit of large *N > P* and for random patterns, **P** becomes semi-orthogonal and left-invertible, turning the pseudoinverse in Eq. 29 into a transpose.

To find the neuronal forgetting times *τ*_*i*_ from Eq. 1, we extract the diagonal entries of the recurrent dynamics in neural space **A**^rec^ and interpret them as the effective reciprocal neuronal time constants 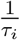 for each neuron *i*:

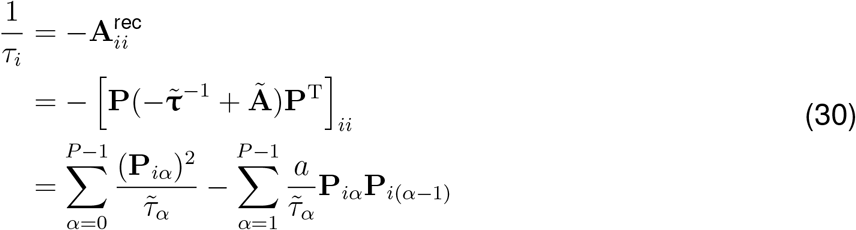

Due to the central limit theorem, the inverse neuronal forgetting times 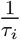 are approximately normally distributed for a sufficiently large pattern number *P* . Their expectation value and variance depend on the statistics of the pattern vector entries **P**_*iα*_ which we assume to be independent, Gaussian random variables with zero mean and finite variance, i.e.

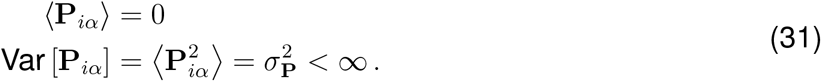

To compute the mean and variance of the inverse neuronal forgetting times, we need to compute the mean and variance of the two pattern-dependent terms in Eq. 30:

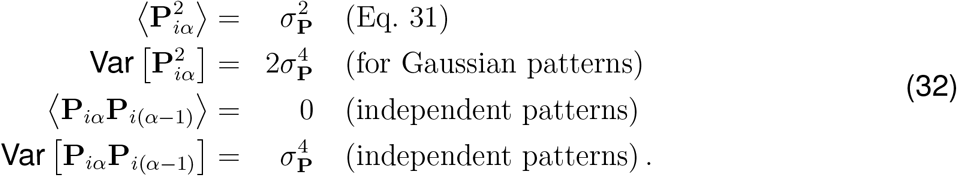

The mean of the inverse neuronal forgetting time is then given by:

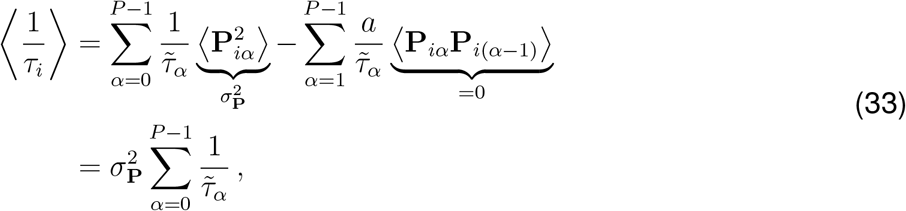

and its variance is:

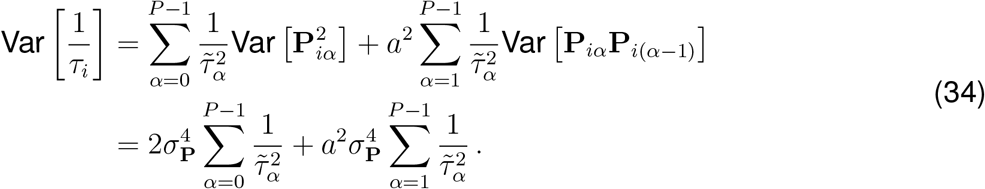

To keep the scale of the neuronal representation/engrams independent of the number of patterns, the patterns must be normalized by the square root of their number. In the simulations, we chose

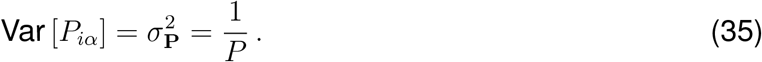

Inserting this into the expressions for the mean and the variance of the inverse neuronal time constants yields:

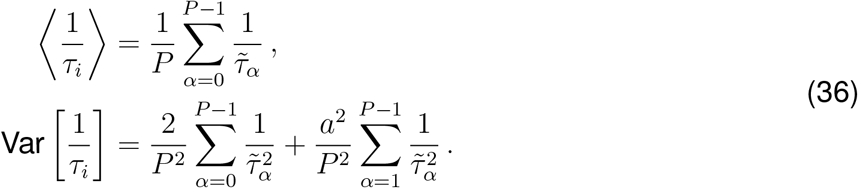

Because the forgetting times of the patterns are geometrically distributed (Eq. 28), these sums are geometric series and can be solved analytically:

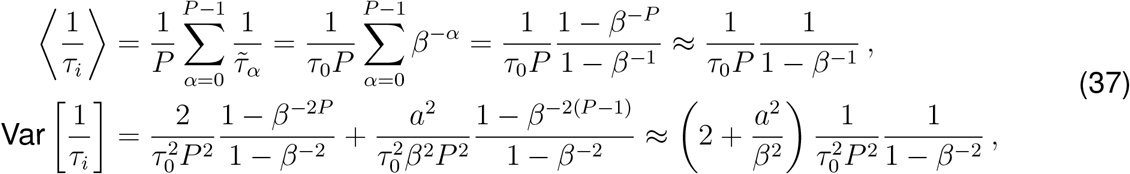

where the approximations become accurate for a large pattern number *P* .

Based on these expressions, we can compute the coefficient of variation of the inverse time constants as a measure of relative variability:

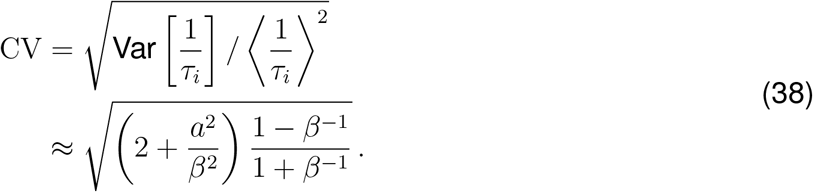

For the parameter choices in the model (*a* = 1, *P* = 500, *β* = 10000^1*/*(*P* −1)^), this yields a CV of approximately 0.165. The inverse neuronal time constants are therefore relatively homogeneous. The same is true for the time constants themselves, although in a strict sense, the mean and variance of the inverse of a Gaussian variable do not exist.

The neuronal forgetting times *τ*_*i*_ are the reciprocal of the normally distributed variable 1*/τ*_*i*_, and thus follow a reciprocal normal distribution with probability density function

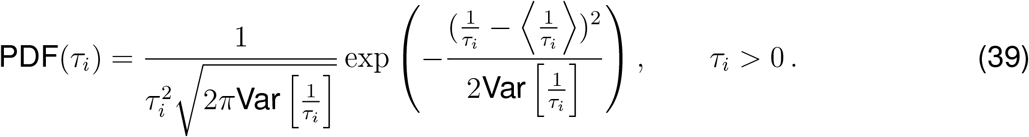

### S4 Effects of subsampling for correlated stimulus tuning

**Figure S4.1:**
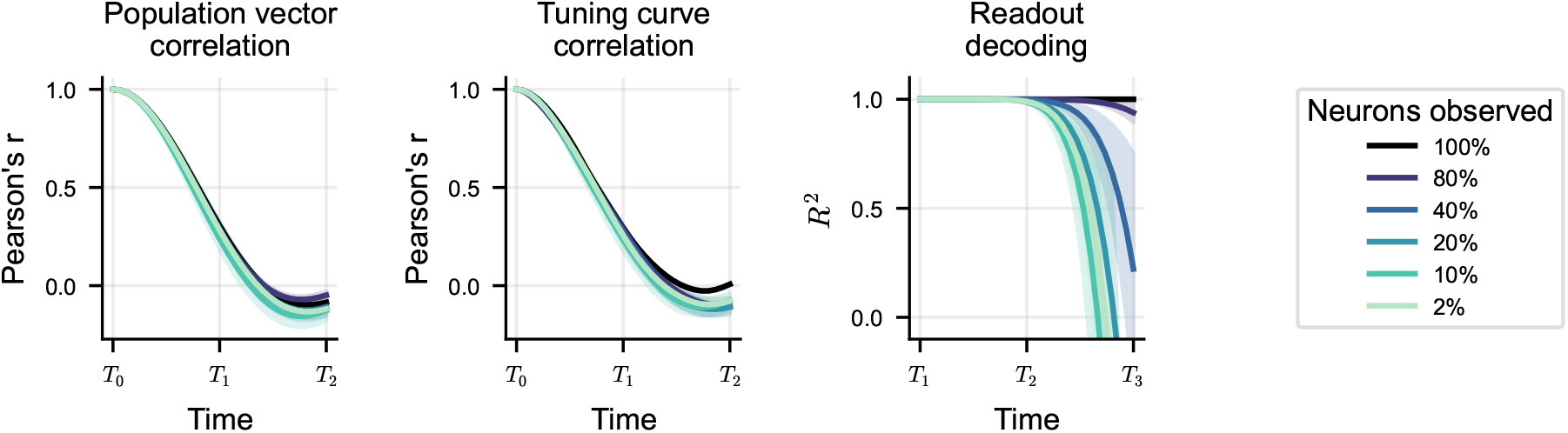
For correlated stimulus tuning (same simulations as shown in Fig. 5I,J), the results on population vector correlation, tuning curve correlation and readout decoding are qualitatively similar to those for random stimulus tuning (Fig. 5D-F).

### S5 Mathematical derivation of apparent stochasticity under subsampling

To characterize the effect of unrecorded neuronal activity on the recorded engram dynamic, we model the two corresponding neural populations as interacting physical systems: a measured subsystem coupled to a bath. For analytical tractability we restrict our calculations to the case of rotational consolidation dynamics and exclude the forgetting component.

Using Green’s function formalism we calculate measurable properties of the engram dynamics, such as the cosine similarity between one engram at two moments in time, an analogous to a time-autocorrelation function. We quantify the contributions from the coupling with the bath using self-energy calculations and we derive the explicit dependence on the sampling fraction (the fraction of neurons that are observed). We derive an exact solution for the time-autocorrelation in the case of Gaussian random hamiltonians in the thermodynamic limit. We show that the coupling with the bath causes a decay in the time-autocorrelation of rotation eigenstates, indicating that the unobserved population effectively acts as noise on the observed subspace. Analytical results are compared with numerical simulations that show a qualitative agreement already in systems with *N* = 10^3^ neurons.

These calculation indicate that, even with a deterministic model, engram dynamics might appear stochastic if observed on a subset of neurons.

#### Full engram dynamics

The full dynamics for an engram **e** is given by

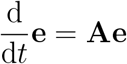

To generate rotations, the *N* × *N* consolidation matrix **A** must be taken anti-symmetric: **A**_*ij*_ = −**A**_*ji*_. Multiplying the equation above by the imaginary unit *i*, one obtains an analogous of the Schrödinger equation, with the Hermitian Hamiltonian **H** = *i***A**:

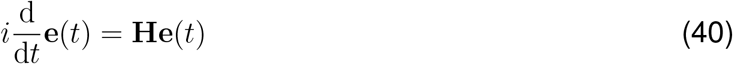

Let *E*_*i*_ be the eigenvalues of the Hamiltonian^1^ and ***ψ***_*i*_ its eigenvectors, which form a basis for the space of engrams: 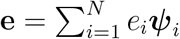. The solution to the engram dynamical model Eq. 40 takes a simple form in this basis:

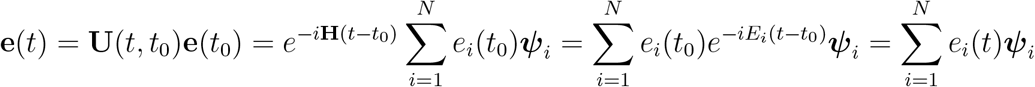

We introduced the time-evolution operator **U**(*t, t*_0_) = exp[− *i***H**(*t* − *t*_0_)]. The dynamics for an eigenstate ***ψ***_*i*_ of the consolidation Hamiltonian is given by a rotation with frequency 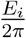:

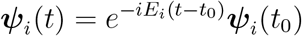

One common mathematical tool to study interacting systems are Green’s functions. We define the *Green’s function* operator **G**(*t, t*_0_) as the solution to the following equation:

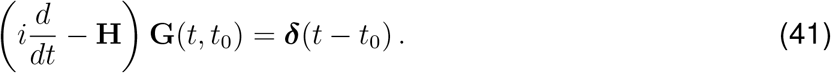

Green’s functions are generally not unique. The one of our specific interest is called *retarded Green’s function*, which connects two points in time that are causally related (*t > t*^*′*^). In the temporal and the Fourier domain, it reads

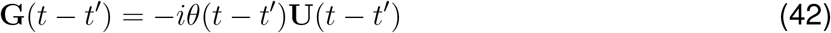

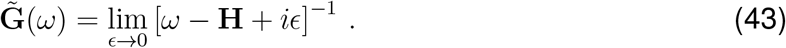

The matrix elements for one eigenstate ***ψ***_*j*_ are

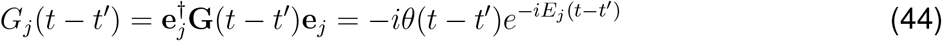

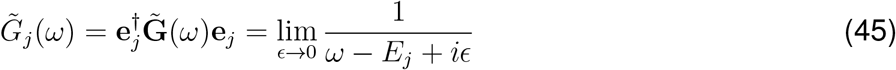

The Green’s function for a generic engram (in time or frequency domain) is

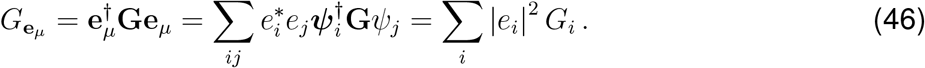

Green’s functions are tightly related to temporal correlations: the cosine similarity 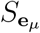 between **e**_*µ*_ and itself at later time *t >* 0 is given by

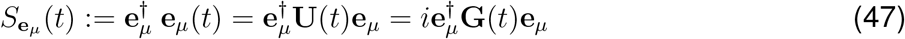

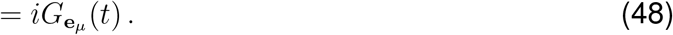

For real engrams, the cosine similarity is a real number, and it can be expressed in terms of the spectral function 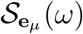 via Fourier transform:

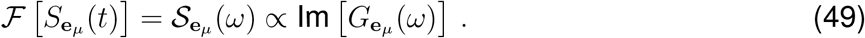

#### Gaussian random Hamiltonians

We now consider Hamiltonians **H** = *i***A** to be Gaussian random *N* × *N* matrices, with purely imaginary elements **H**_*ij*_ = *i***A**_*ij*_. Above-diagonal entries **A**_*i<j*_ are independent random variables with identical normal distribution 𝒩 (0, *σ*_*H*_). Below-diagonal elements, instead, are fixed by antisymmetry: **A**_*ij*_ = −**A**_*ji*_. To properly scale the transformation generated by the Hamiltonian with the number of neurons *N*, we set 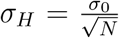. In the continuum limit (*N* → ∞), the distribution of the eigenvalues *E*_*i*_ converges to the semicircular distribution (Fig. S5.1A)

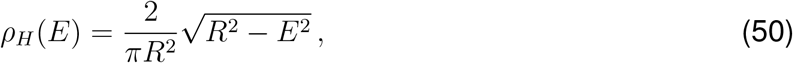

defined in the interval 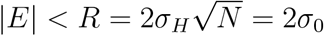. For each Gaussian random Hamiltonian, we can apply the results of Equations 44 and 45 to compute the Green’s function of an **H**-eigenstate ***ψ***_*i*_ with eigenvalue *E*_*i*_. As before, the time-autocorrelation will oscillate in time with frequency 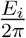, without decaying.

In general, we might assume an engram to be a random superposition of eigenstates of the rotational consolidation matrix:

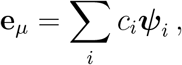

with *c*_*i*_ randomly sampled from a distribution with zero mean, and standard deviation 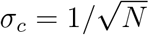. Its average Green’s function is given by

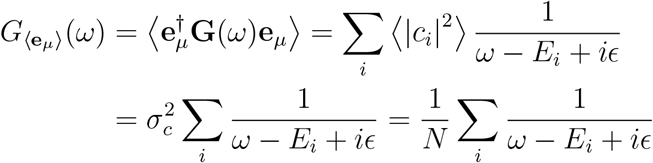

For large *N* we use the continuous approximation:

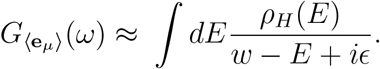

This integral corresponds to the Stieltjes transformation of the semicircle law. It can be solved performing a geometric series expansion of *ρ*_*H*_(*E*), and interchanging summation and integration. Alternative solutions make use of analytical continuation and contour integration. We refer to Jiang (2021) and to Bai and Silverstein (2010) for detailed calculations. Instead, we report here an intuitive derivation of the result. Multiplying both numerator and denominator by (*ω* − *E* + *iϵ*), we obtain

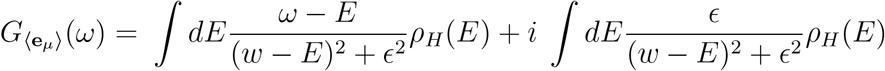

We recall one definition for the Dirac-*δ* in terms of Lorentzian distributions 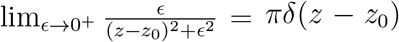. The imaginary part of the integral, which relates to the spectral function 𝒮 (*ω*), is therefore given by the density of eigenvalues:

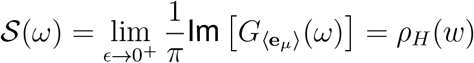

We can therefore reconstruct the autocorrelation function for a random state **e**_*µ*_ by inverse Fourier transform of the semicircle law:

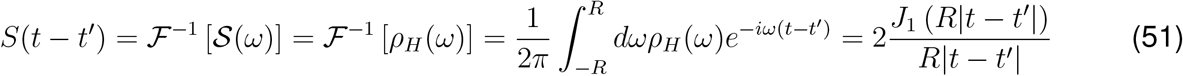

Here, *J*_1_(*x*) is a Bessel function of the first kind of order 1. The autocorrelation *S*(*t*) presents a decaying oscillation (Figure S5.1). For large *t*, the decay is described by a power law *S*(*t*) ∼ *t*^−3*/*2^.

**Figure S5.1:**
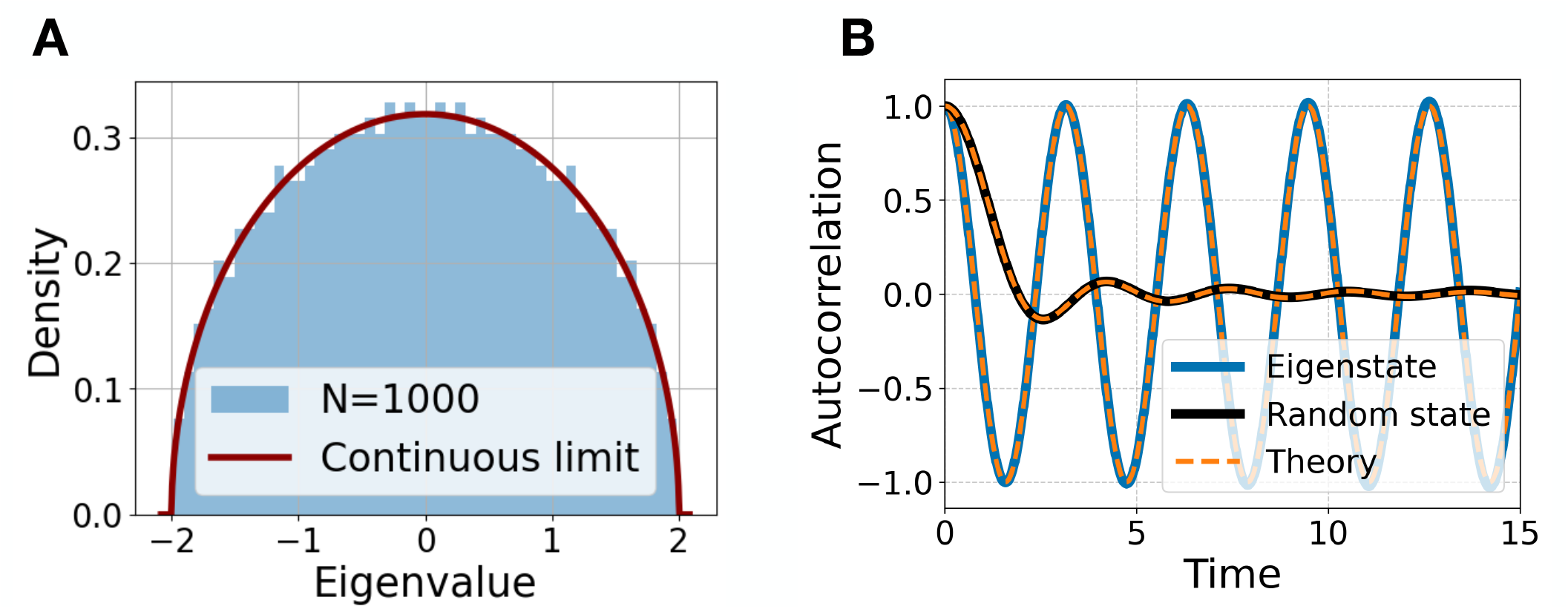
Dynamics in the fully-observed system. **(A)**The distribution of the eigenvalues for Gaussian random matrix **H** with size *N* = 1000 and entries standard deviation 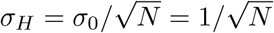. The red line corresponds to the semicircle law Eq.50. **(B)** Time-autocorrelation: eigenstate of the rotation hamiltonian (blue line) and average autocorrelation of 50 independent random states (black line). The orange lines corresponds to the analytical expressions in Eq. 45 and Eq. 51.

Although the dynamics is deterministic and purely rotational, the autocorrelation of a generic (random) state decays in time. A generic engram evolves in time becoming, on average, uncorrelated to its initial representation, as shown in Fig. 4D. This result might appear counterintuitive when picturing rotations in low dimensional spaces. However, rotations in high-dimensions can be expressed as a composition of several independent planar rotations with incommensurate frequencies (the eigenvalues ±*E*_*i*_). This random superposition of independently-rotating modes (the eigenstates ***ψ***_*i*_) generates a complex trajectory, in which the engram takes a long time (infinite, in the limit of *N* → ∞) to return to its initial state.

#### The problem of subsampling

The goal of this section is to compute the effect of subsampling, that is, how the rotational dynamics is changed when only a subspace of the total engram space is observed (for instance, measuring a fraction of the total number of neurons). We model the observed and unobserved subspaces as two coupled physical systems. We referred to the former as the “observed” or “sampled” system, and to the latter as a “bath” (in analogy to the physics of open systems). The full Hamiltonian **H** is rewritten as a composition of the following blocks:

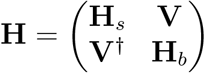

Here, **H**_*s*_ and **H**_*b*_ are the Hamiltonians of the observed subsystem and the bath, with dimensions *N*_*s*_ and *N*_*b*_ respectively. The interaction between the two systems is described by **V** (a *N*_*s*_ × *N*_*b*_ matrix).

We initially apply the Green’s function formalism to the sampled subsystem when the interaction **V** = 0 is momentarily switched off. In this case, the observed dynamics is rotational and purely governed by **H**_*s*_. This Hamiltonian is Hermitian and it can be diagonalized:

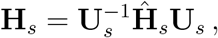

where **Ĥ**_*s*_ is diagonal. Similarly, the dynamics in the bath (the unobserved subspace) is uniquely described by the Hamiltonian 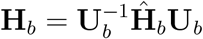. Given an eigenstate 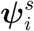 of the sampled subspace with eigenvalue 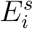, we can use the results of the previous sections to define its respective Green’s function:

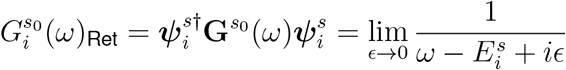

We denote 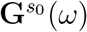 with a superscript *s*_0_ to indicate the absence of coupling (**V** = 0). In case of Gaussian random matrices, the autocorrelation of a random state is given by Eq. 51, with the density of states of the subsystem *ρ*_*s*_(*ω*).

We now switch on the interaction **V** and compute the full Green’s function **G**^*s*^(*ω*), in terms of 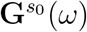 and a correction **Σ**(*ω*), referred to as self-energy. This correction accounts for the coupling with the unobserved neurons, the activity of which affects the dynamics in the observed subspace. The Green’s function in the absence of coupling and the corrected one are commonly referred to as *free* and *interacting* (or *full* ) propagators.

#### Self-energy formalism

To compute the full propagator **G**^*s*^ we start with the definition of the Green’s function operator in Fourier space:

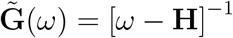

We use the structure of our system to rewrite the relation above as

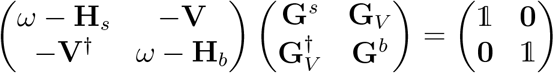

We extract a system of two equations for **G**^*s*^:

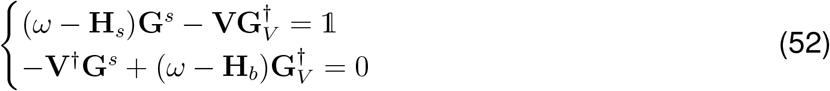

which has the solution

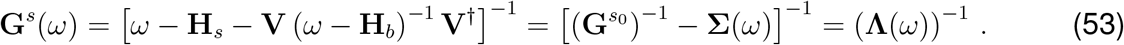

The term **Σ**(*ω*) is commonly named *self-energy* and it describes the correction to the free propagator 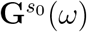, due to the coupling with the bath. This expression leads to a self-consistent relation for **G**^*s*^(*ω*), known as the Dyson equation:

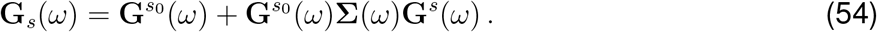

We can express Eq. 53 in the diagonal representation, diagonalizing both the Hamiltonian of the observed subsystem and the Hamiltonian of the bath:

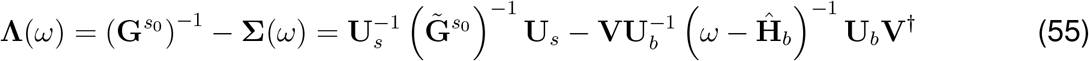

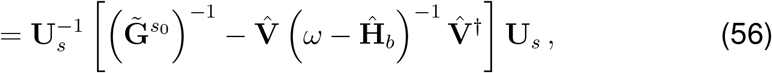

where we defined 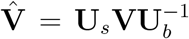. We compute now the explicit expression for the matrix elements of **Λ**(*ω*) in the diagonal representation: 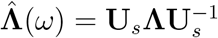. The diagonal elements read:

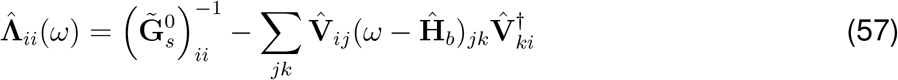

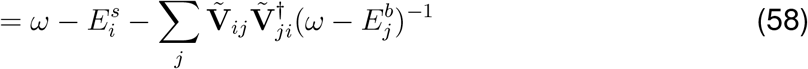

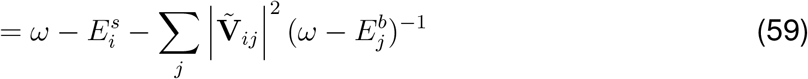

Similarly, the off-diagonal elements are given by

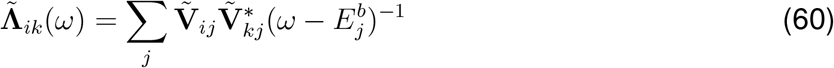

The full Green’s function operator is given by the inverse of the matrix **Λ**, which is generally non-diagonal. In the next section we focus on the case of Gaussian random matrix, for which this result can be simplified.

#### Self-energy for Gaussian random matrices

We are interested now in the case of the full Hamiltonian **H** being a Gaussian random matrix, with independent entries that are normally distributed: 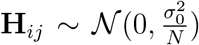. The results of the previous section can be simplified by an ensemble average of the matrix elements of 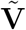 (assuming a self-averaging in this large neuronal system):

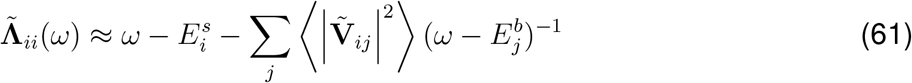

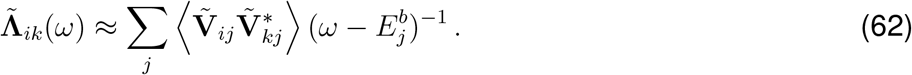

The two averages read

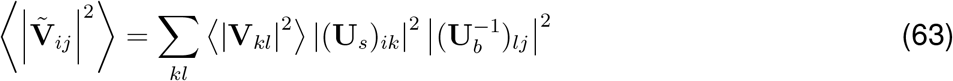

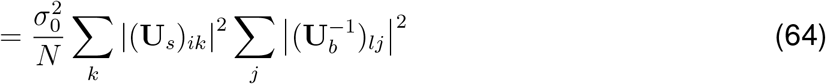

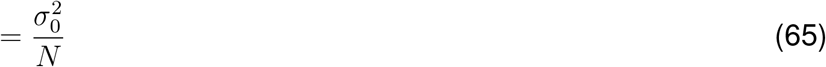

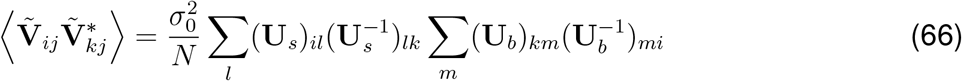

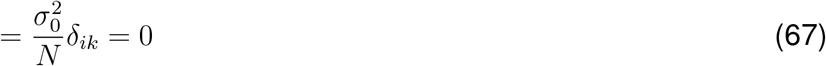

where we used the orthogonality of **U**_*s*_ and **U**_*b*_. The elements of the matrix **Λ**(*ω*) read:

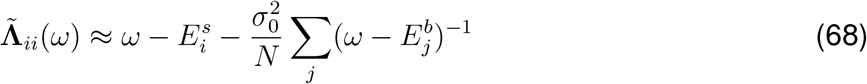

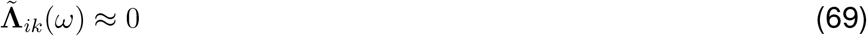

We can recover the full Green’s function for an eigenstate 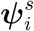 of the observed subsystem from the diagonal components of 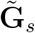:

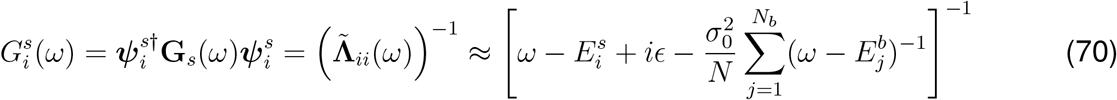

The self-energy for Gaussian random matrices is given by the sum of contributions from all the modes of the bath 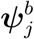, which energy is 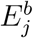:

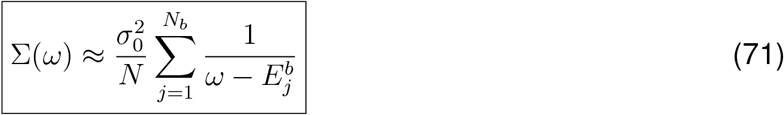

#### Thermodynamic limit

We now focus on the case for large number of dimensions *N*, which can be studied in the continuum approximation where *N* → ∞. In this setting, the result of the previous section becomes exact. Because the system is split into two parts (*N* = *N*_*s*_ + *N*_*b*_) one can make different continuum approximations by taking different continuous limits in the two subspaces. Whether one sends to infinity *N*_*s*_ or *N*_*b*_ or both - and if both, at what relative speed - determines different thermodynamic limits. In our model, we are concerned with the problem of observing only a fraction of the total number of neurons and how the observed dynamics depend on this fraction. It is therefore natural to consider the thermodynamic limit in which the fraction is preserved. This is obtained by sending the size of both the sampled subsystem and the bath to infinity, while keeping their fraction fixed:

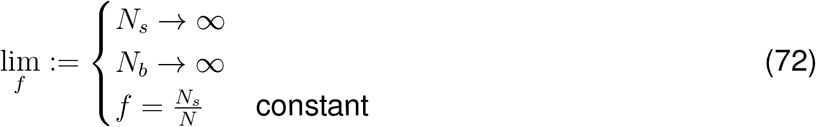

In the absence of coupling, the distribution of eigenvalues in the sampled subsystem and in the bath are described by semicircle laws:

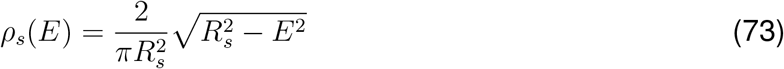

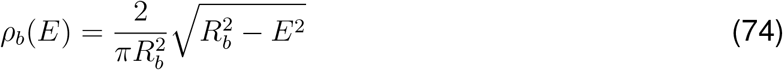

for |*E*| *< R*_*s/b*_. The radii of the two semicircles are given by

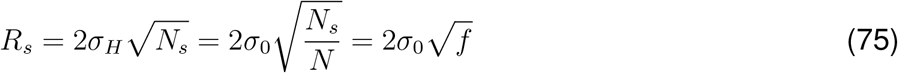

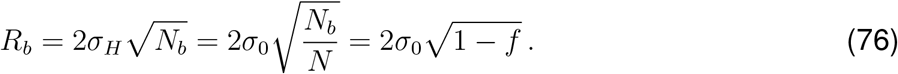

The continuum limit allows us to simplify the self-energy Eq. 71, by transforming the summation over the modes of the bath into an integral weighted by the density of eigenstates. Specifically

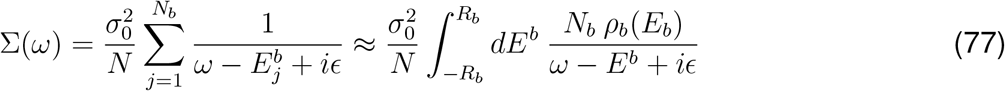

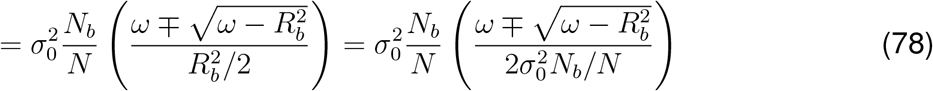

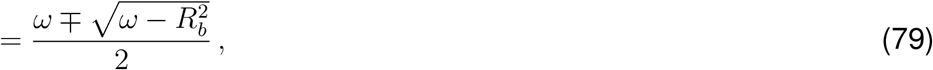

where the ∓ is taken such that 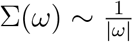 for *ω* → ∞. The integral above is the Stieltjes transform of the semicircle law^2^ We separate the self-energy into its real and imaginary parts:

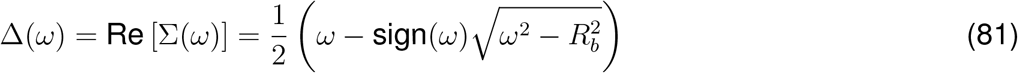

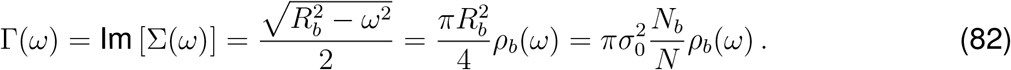

Note that the second term in Δ(*ω*) vanishes for |*ω*| *< R*_*b*_, and viceversa the imaginary part Γ(*ω*) vanishes for |*ω*| *> R*_*b*_. The full Green’s function takes the following form:

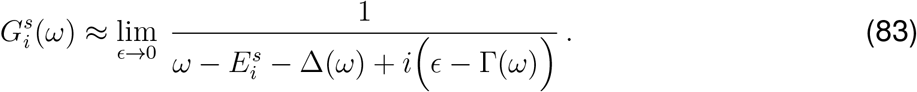

The spectral function is computed as

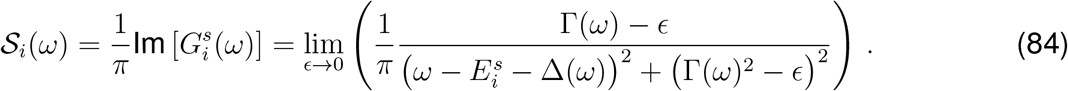

We study this limit in two different cases, namely for Γ(*ω*) ≠ 0 and Γ(*ω*) = 0. These are given by |*ω*| *< R*_*b*_ and |*ω*| *> R*_*b*_ respectively, and are indicated by _*<*_ and _*>*_ subscripts. In the first case, the spectral function reads:

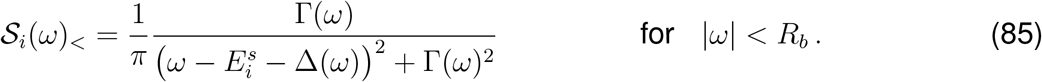

In the second case instead, the function reads

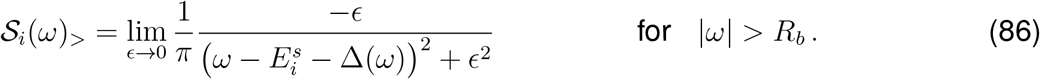

The behavior of this second limit depends on the zero of the term 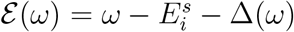, which corresponds to shifted energy of the original mode^3^. If ℰ (*ω*) vanishes at |*ω*| *> R*_*b*_, the limit in Eq.86 gives a finite contribution. If, instead, the zero of ℰ (*ω*) lies in [−*R*_*b*_, *R*_*b*_], the limit for 𝒮_*i*_(*ω*)_*>*_ is zero, and the only contribution comes from 𝒮_*i*_(*ω*)_*<*_. This second case is always given when 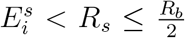, that corresponds to 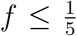. We refer to this situation as “low sampling fraction”. Figure S5.2A shows how theory captures the behavior of the autocorrelation and spectral density. If instead 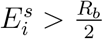, the limit reads

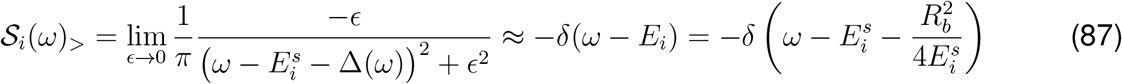

The spectral function, which was peaked at one frequency when no coupling was present, takes now finite values at all frequencies |*ω*| *< R*_*b*_. This is a consequence of the coupling with the bath, and it results in a decay of time-autocorrelations. This effect, known as *damping* or *dephasing*, gives a finite life-time to the rotational eigenstates in the observed subspace. An analogous damping profile would be observed in the presence of stochasticity in the engram dynamics. The similarity is not surprising, since the coupling with the bath can be interpreted as a noise source in the dynamics of the sampled subsystem.

In conclusion, the autocorrelation of a rotational eigenstates, that would oscillate in a fully-observed system, decays in the presence of subsampling, because of the coupling with the unobserved population. In the case of *low* sampling fraction, this dephasing is equivalent to the autocorrelation decay in the full system for a random engram (Eq. 51). This is not surprising, since the bath – whose contribution on the dynamics dominates for low sampling fraction – has the same statistics as the full system. However, the substantial difference between the dephasing caused by subsampling (Fig. S5.2B-D) and the decay in the fully observed system for a random engram (Fig. S5.1B), is that in the latter the dynamics is predictable. This can be seen in the linear dynamical system fits to simulated engram dynamics, where in the fully observed system the fitting succeeds while it fails in a partially observed system (Fig. 5I). This analysis shows that characterizing and fitting engram dynamics from data is in general strongly limited by the problem of subsampling.

#### Numerical simulations

We simulate the rotational dynamics sampling an *N* × *N* anti-symmetric matrix **H**. Above diagonal entries **H**_*i<j*_ are independent random variables with identical normal distribution 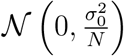. All the remaining entries are fixed by anti-symmetry. We simulate the decorrelation of one eigenstate 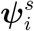 of **H**_*s*_ with energy 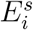 (that is, one rotational eigenstate of the sampled subsystem), caused by the coupling with the bath. We define the initial state as following: the first *N*_*s*_ dimensions correspond to ***ψ***_*i*_, and the remaining *N*_*b*_ are set to zero to reflect no initial activity in the bath: **e**(0) = (***ψ***_*i*_ **0**_*b*_ )^*T*^ . Here, we referred to **0** as a zero-vector of *N* components. The dynamics is simulated using Euler integration, and the time-autocorrelation is measured by measuring at the activity **e**_*s*_(*t*) in the first *N*_*s*_ dimensions: 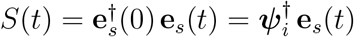.

**Figure S5.2:**
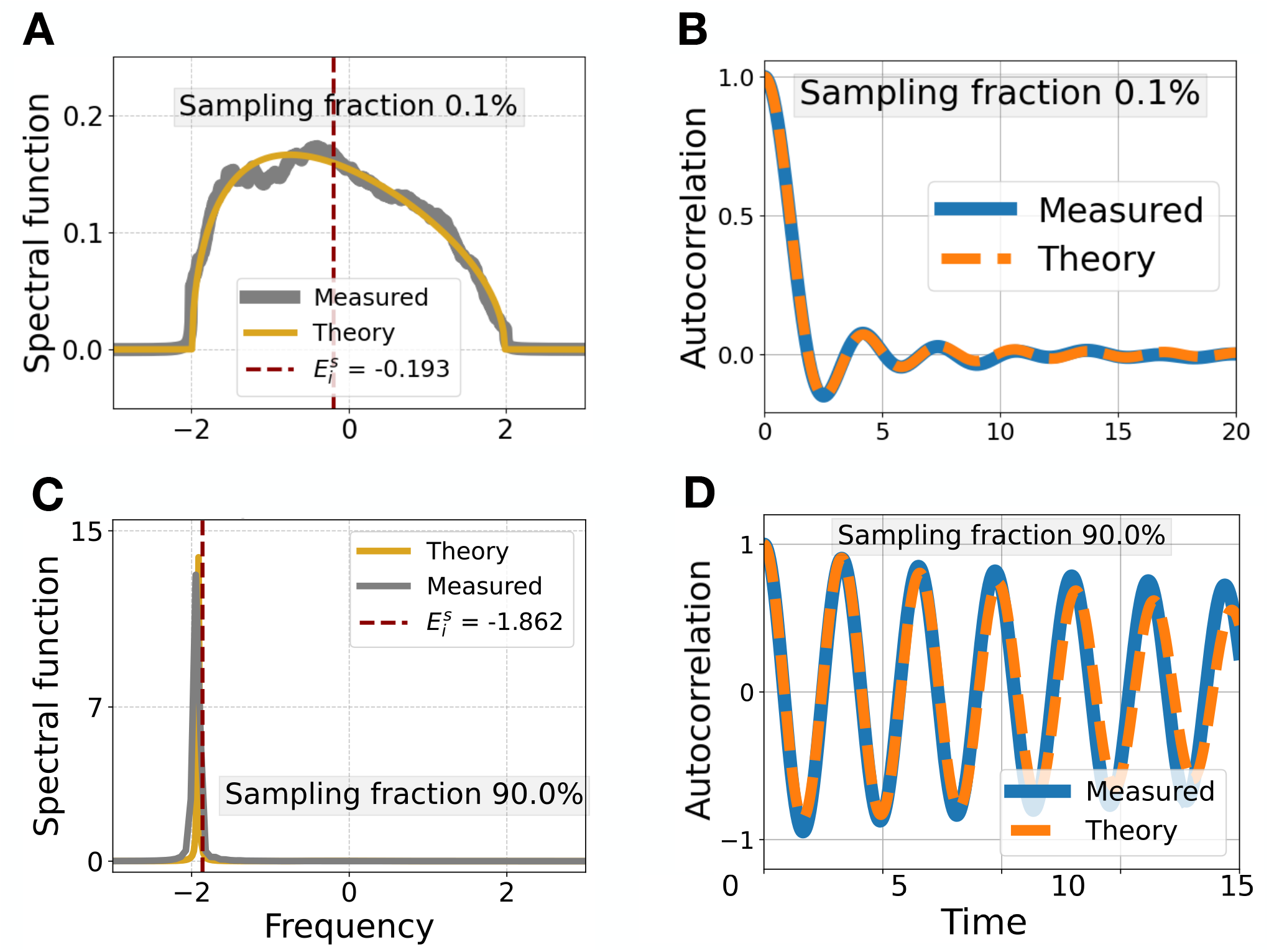
**Low and high sampling fractions** Spectral functions (**A**,**C**) and time-autocorrelations (**B**,**D**) of a rotation eigenstate 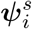 of the sampled sub-system, with energy 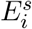. Numerical values are obtained simulating a full rotation hamiltonian **H** of size *N* = 10000, using Euler integration (time constant *τ* = 1, time step Δ*t* = 0.001, simulation time length *T*_*f*_ = 1000). The activity is observed in the first *N*_*s*_ = 100 (**A, B**) and *N*_*s*_ = 9000 (**C, D**) dimensions. *σ*_0_ = 1.

## Footnotes

1 The anti-symmetric structure of **H** implies that for each eigenvalue *E*_*i*_ ∈ ℝ, there is one of opposite sign −*E*_*i*_.

2 Given the semicircle distribution 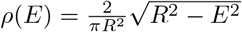, the Stieltjes transform reads 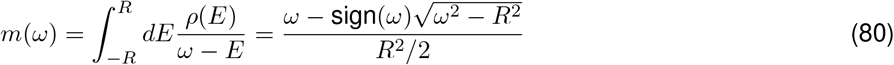

3 Since the function ℰ (*ω*) is continuous, monotonic and lim_*ω*→*±*∞_ ℰ (*ω*) = ±∞, it has only one zero.

## REFERENCES

Benna, M.K., and Fusi, S. (2016). Computational principles of synaptic memory consolidation. Nature Neuroscience 19, 1697–1706. doi: 10.1038/nn.4401.

Bhasin, B.J., Raymond, J.L., and Goldman, M.S. (2024). Synaptic weight dynamics underlying memory consolidation: Implications for learning rules, circuit organization, and circuit function. Proceedings of the National Academy of Sciences 121, e2406010121. doi: 10.1073/pnas.2406010121.

Cervantes-Sandoval, I., Martin-Peña, A., Berry, J.A., and Davis, R.L. (2013). System-like consolidation of olfactory memories in Drosophila. Journal of Neuroscience 33, 9846–54. doi: 10.1523/JNEUROSCI.0451-13.2013.

Chouhan, N.S., Griffith, L.C., Haynes, P., and Sehgal, A. (2020). Availability of food determines the need for sleep in memory consolidation. Nature 589, 582–585. doi: 10.1038/s41586-020-2997-y.

Clopath, C. (2012). Synaptic consolidation: An approach to long-term learning. Cognitive Neurodynamics 6, 251–257. doi: 10.1007/s11571-011-9177-6.

de Snoo, M.L., and Frankland, P.W. (2025). Neurobiological mechanisms of forgetting across timescales. Current Opinion in Neurobiology 90, 102972. doi: 10.1016/j.conb.2025.102972.

Debiec, J., LeDoux, J.E., and Nader, K. (2002). Cellular and Systems Reconsolidation in the Hippocampus. Neuron 36, 527–538. doi: 10.1016/S0896-6273(02)01001-2.

Deitch, D., Rubin, A., and Ziv, Y. (2021). Representational drift in the mouse visual cortex. Current Biology 31, 4327–4339.e6. doi: 10.1016/j.cub.2021.07.062.

Delamare, G., Zaki, Y., Cai, D.J., and Clopath, C. (2024). Drift of neural ensembles driven by slow fluctuations of intrinsic excitability. eLife 12, RP88053. doi: 10.7554/eLife.88053.

Devalle, F., Zou, L., Cecchini, G., and Roxin, A. (2025). Representational drift as the consequence of ongoing memory storage. Scientific Reports 15, 27746. doi: 10.1038/s41598-025-11102-x.

Diekelmann, S., and Born, J. (2010). The memory function of sleep. Nature Reviews Neuroscience 11, 114–126. doi: 10.1038/nrn2762.

Driscoll, L.N., Duncker, L., and Harvey, C.D. (2022). Representational drift: Emerging theories for continual learning and experimental future directions. Current Opinion in Neurobiology 76, 102609. doi: 10.1016/j.conb.2022.102609.

Driscoll, L.N., Pettit, N.L., Minderer, M., Chettih, S.N., and Harvey, C.D. (2017). Dynamic Reorganization of Neuronal Activity Patterns in Parietal Cortex. Cell 170, 986–999.e16. doi: 10.1016/j.cell.2017.07.021.

Eppler, J.B., Lai, T., Aschauer, D.F., Rumpel, S., and Kaschube, M. (2026). Representational drift reflects ongoing balancing of stochastic changes by Hebbian learning. Proceedings of the National Academy of Sciences 123, e2503046123. doi: 10.1073/pnas.2503046123.

Fusi, S. (2017). Computational models of long term plasticity and memory. Peprint at arXiv. doi: 10.48550/arXiv.1706.04946.

Fusi, S., Drew, P.J., and Abbott, L. (2005). Cascade Models of Synaptically Stored Memories. Neuron 45, 599–611. doi: 10.1016/j.neuron.2005.02.001.

Ganguli, S., Huh, D., and Sompolinsky, H. (2008). Memory traces in dynamical systems. Proceedings of the National Academy of Sciences 105, 18970–18975. doi: 10.1073/pnas.0804451105.

Geva, N., Deitch, D., Rubin, A., and Ziv, Y. (2023). Time and experience differentially affect distinct aspects of hippocampal representational drift. Neuron 111, 2357–2366.e5. doi: 10.1016/j.neuron.2023.05.005.

Golbabaei, A., Coelho, C.A.O., de Snoo, M.L., De Cristofaro, A., Josselyn, S.A., and Frankland, P.W. (2025a). Neurogenesis-dependent transformation of hippocampal memory traces during systems consolidation. Current Biology 35, 4959–4969.e4. doi: 10.1016/j.cub.2025.09.005.

Golbabaei, A., Josselyn, S.A., and Frankland, P.W. (2025b). PV-dependent reorganization of prelimbic cortex sub-engrams during systems consolidation. Neuron. doi: 10.1016/j.neuron.2025.09.033.

Goldman, M.S. (2009). Memory without Feedback in a Neural Network. Neuron 61, 621–634. doi: 10.1016/j.neuron.2008.12.012.

Haimerl, C., and Machens, C. (2025). Representational drift without synaptic plasticity. Preprint at bioRxiv. doi: 10.1101/2025.07.23.666352.

Han, J.H., Kushner, S.A., Yiu, A.P., Hsiang, H.L.L., Buch, T., Waisman, A., Bontempi, B., Neve, R.L., Frankland, P.W., and Josselyn, S.A. (2009). Selective Erasure of a Fear Memory. Science 323, 1492–1496. doi: 10.1126/science.1164139.

Himmer, L., Schönauer, M., Heib, D.P.J., Schabus, M., and Gais, S. (2019). Rehearsal initiates systems memory consolidation, sleep makes it last. Science Advances 5, eaav1695. doi: 10.1126/sciadv.aav1695.

Hopfield, J.J. (1982). Neural networks and physical systems with emergent collective computational abilities. Proceedings of the National Academy of Sciences 79, 2554–8. doi: 10.1073/PNAS.79.8.2554.

Josselyn, S.A., and Tonegawa, S. (2020). Memory engrams: Recalling the past and imagining the future. Science 367. doi: 10.1126/science.aaw4325.

Kalle Kossio, Y.F., Goedeke, S., Klos, C., and Memmesheimer, R.M. (2021). Drifting assemblies for persistent memory: Neuron transitions and unsupervised compensation. Proceedings of the National Academy of Sciences 118, e2023832118. doi: 10.1073/pnas.2023832118.

Kaufman, M.T., Churchland, M.M., Ryu, S.I., and Shenoy, K.V. (2014). Cortical activity in the null space: permitting preparation without movement. Nature neuroscience 17, 440–448. doi: 10.1038/nn.3643.

Khatib, D., Ratzon, A., Sellevoll, M., Barak, O., Morris, G., and Derdikman, D. (2023). Active experience, not time, determines within-day representational drift in dorsal CA1. Neuron 111, 2348–2356.e5. doi: 10.1016/j.neuron.2023.05.014.

Kossio, F.Y.K., and Memmesheimer, R.M. (2025). Brain-wide representational drift: Memory consolidation and entropic force. Preprint at bioRxiv. doi: 10.1101/2025.09.25.678547.

Lee, J.H., Kim, W.B., Park, E.H., and Cho, J.H. (2023). Neocortical synaptic engrams for remote contextual memories. Nature Neuroscience 26, 259–273. doi: 10.1038/s41593-022-01223-1.

Levina, A., Priesemann, V., and Zierenberg, J. (2022). Tackling the subsampling problem to infer collective properties from limited data. Nature Reviews Physics 4, 770–784. doi: 10.1038/s42254-022-00532-5.

Liu, X., Ramirez, S., Pang, P.T., Puryear, C.B., Govindarajan, A., Deisseroth, K., and Tonegawa, S. (2012). Optogenetic stimulation of a hippocampal engram activates fear memory recall. Nature 484, 381–385. doi: 10.1038/nature11028.

Martin, S., and Morris, R. (2002). New life in an old idea: The synaptic plasticity and memory hypothesis revisited. Hippocampus 12, 609–636. doi: 10.1002/hipo.10107.

McClelland, J.L., McNaughton, B.L., and O’Reilly, R.C. (1995). Why there are complementary learning systems in the hippocampus and neocortex: Insights from the successes and failures of connectionist models of learning and memory. Psychological review 102, 419–57. doi: 10.1037/0033-295X.102.3.419.

Micou, C., and O’Leary, T. (2023). Representational drift as a window into neural and behavioural plasticity. Current Opinion in Neurobiology 81, 102746. doi: 10.1016/j.conb.2023.102746.

Micou, C., and O’Leary, T. (2024). Heavy-tailed statistics of cortical representational drift are advantageous for stabilised downstream readouts. Preprint at bioRxiv. doi: 10.1101/2024.09.26.614914.

Morales, G.B., Muñoz, M.A., and Tu, Y. (2024). Representational Drift and Learning-Induced Stabilization in the Olfactory Cortex. Peprint at arXiv. doi: 10.48550/arXiv.2412.13713.

Moscovitch, M., and Gilboa, A. (2022). Has the concept of systems consolidation outlived its use-fulness? Identification and evaluation of premises underlying systems consolidation. Faculty Reviews 11, 33. doi: 10.12703/r/11-33.

Nadel, L., and Moscovitch, M. (1997). Memory consolidation, retrograde amnesia and the hippocampal complex. Current Opinion in Neurobiology 7, 217–227. doi: 10.1016/S0959-4388(97)80010-4.

Natrajan, M., and Fitzgerald, J.E. (2024). Stability through plasticity: Finding robust memories through representational drift. Preprint at bioRxiv. doi: 10.1101/2024.12.19.629245.

Owald, D., and Waddell, S. (2015). Olfactory learning skews mushroom body output pathways to steer behavioral choice in Drosophila. Current Opinion in Neurobiology 35, 178–184. doi: 10.1016/J.CONB.2015.10.002.

Qian, W., Zavatone-Veth, J.A., Ruben, B.S., and Pehlevan, C. (2024). Partial observation can induce mechanistic mismatches in data-constrained models of neural dynamics. In The Thirty-eighth Annual Conference on Neural Information Processing Systems.

Qin, S., Farashahi, S., Lipshutz, D., Sengupta, A.M., Chklovskii, D.B., and Pehlevan, C. (2023). Coordinated drift of receptive fields in Hebbian/anti-Hebbian network models during noisy representation learning. Nature Neuroscience 26, 339–349. doi: 10.1038/s41593-022-01225-z.

Ratzon, A., Derdikman, D., and Barak, O. (2024). Representational drift as a result of implicit regularization. eLife 12. doi: 10.7554/eLife.90069.2.

Remme, M.W.H., Bergmann, U., Alevi, D., Schreiber, S., Sprekeler, H., and Kempter, R. (2021). Hebbian plasticity in parallel synaptic pathways: A circuit mechanism for systems memory consolidation. PLOS Computational Biology 17, e1009681. doi: 10.1371/journal.pcbi.1009681.

Reymann, K.G., and Frey, J.U. (2007). The late maintenance of hippocampal LTP: Requirements, phases, ‘synaptic tagging’, ‘late-associativity’ and implications. Neuropharmacology 52, 24– 40. doi: 10.1016/j.neuropharm.2006.07.026.

Robertson, E.M. (2012). New Insights in Human Memory Interference and Consolidation. Current Biology 22, R66–R71. doi: 10.1016/j.cub.2011.11.051.

Roxin, A., and Fusi, S. (2013). Efficient Partitioning of Memory Systems and Its Importance for Memory Consolidation. PLoS Computational Biology 9, e1003146. doi: 10.1371/journal.pcbi.1003146.

Roy, D.S., Park, Y.G., Kim, M.E., Zhang, Y., Ogawa, S.K., DiNapoli, N., Gu, X., Cho, J.H., Choi, H., Kamentsky, L., Martin, J., Mosto, O., Aida, T., Chung, K., and Tonegawa, S. (2022). Brain-wide mapping reveals that engrams for a single memory are distributed across multiple brain regions. Nature Communications 13, 1799. doi: 10.1038/s41467-022-29384-4.

Rule, M.E., Loback, A.R., Raman, D.V., Driscoll, L.N., Harvey, C.D., and O’Leary, T. (2020). Stable task information from an unstable neural population. eLife 9, e51121. doi: 10.7554/eLife.51121.

Rule, M.E., and O’Leary, T. (2022). Self-healing codes: How stable neural populations can track continually reconfiguring neural representations. Proceedings of the National Academy of Sciences 119, e2106692119. doi: 10.1073/pnas.2106692119.

Rule, M.E., O’Leary, T., and Harvey, C.D. (2019). Causes and consequences of representational drift. Current Opinion in Neurobiology 58, 141–147. doi: 10.1016/j.conb.2019.08.005.

Schlichting, M.L., Mumford, J.A., and Preston, A.R. (2015). Learning-related representational changes reveal dissociable integration and separation signatures in the hippocampus and prefrontal cortex. Nature Communications 6, 8151. doi: 10.1038/ncomms9151.

Schoonover, C.E., Ohashi, S.N., Axel, R., and Fink, A.J.P. (2021). Representational drift in primary olfactory cortex. Nature 594, 541–546. doi: 10.1038/s41586-021-03628-7.

Squire, L.R., and Alvarez, P. (1995). Retrograde amnesia and memory consolidation: A neurobiological perspective. Current Opinion in Neurobiology 5, 169–177. doi: 10.1016/0959-4388(95)80023-9.

Squire, L.R., Genzel, L., Wixted, J.T., and Morris, R.G. (2015). Memory consolidation. Cold Spring Harbor perspectives in biology 7, a021766. doi: 10.1101/cshperspect.a021766.

Sylte, O.C., Kilias, A., Bartos, M., and Sauer, J.F. (2025). Coordinated representational drift supports stable place coding in hippocampal CA1. Preprint at bioRxiv. doi: 10.1101/2025.02.04.636428.

Tomff, D.F., Zhang, Y., Aida, T., Mosto, O., Lu, Y., Chen, M., Sadeh, S., Roy, D.S., and Clopath, C. (2024). Dynamic and selective engrams emerge with memory consolidation. Nature Neuroscience 27, 561–572. doi: 10.1038/s41593-023-01551-w.

Tse, D., Langston, R.F., Kakeyama, M., Bethus, I., Spooner, P.A., Wood, E.R., Witter, M.P., and Morris, R.G.M. (2007). Schemas and memory consolidation. Science 316, 76–82. doi: 10.1126/science.1135935.

Wamsley, E.J. (2022). Offline memory consolidation during waking rest. Nature Reviews Psychology 1, 441–453. doi: 10.1038/s44159-022-00072-w.

Wang, J., Narain, D., Hosseini, E.A., and Jazayeri, M. (2018). Flexible timing by temporal scaling of cortical responses. Nature Neuroscience 21, 102–110. doi: 10.1038/s41593-017-0028-6.

Wilting, J., and Priesemann, V. (2018). Inferring collective dynamical states from widely unobserved systems. Nature Communications 9, 2325. doi: 10.1038/s41467-018-04725-4.

Winocur, G., and Moscovitch, M. (2011). Memory transformation and systems consolidation. Journal of the International Neuropsychological Society 17, 766–780. doi: 10.1017/S1355617711000683.

Winocur, G., Moscovitch, M., and Bontempi, B. (2010). Memory formation and long-term retention in humans and animals: Convergence towards a transformation account of hippocampal–neocortical interactions. Neuropsychologia 48, 2339–2356. doi: 10.1016/J.NEUROPSYCHOLOGIA.2010.04.016.

Wixted, J.T. (2004). The Psychology and Neuroscience of Forgetting. Annual Review of Psychology 55, 235–269. doi: 10.1146/annurev.psych.55.090902.141555.

Zaki, Y., and Cai, D.J. (2024). Memory engram stability and flexibility. Neuropsychopharmacology 50, 285–293. doi: 10.1038/s41386-024-01979-z.

Zhou, Z., and Schapiro, A.C. (2025). A gradient of complementary learning systems emerges through meta-learning. Preprint at bioRxiv. doi: 10.1101/2025.07.10.664201.

Ziv, N.E., and Brenner, N. (2018). Synaptic Tenacity or Lack Thereof: Spontaneous Remodeling of Synapses. Trends in Neurosciences 41, 89–99. doi: 10.1016/j.tins.2017.12.003.

Ziv, Y., Burns, L.D., Cocker, E.D., Hamel, E.O., Ghosh, K.K., Kitch, L.J., Gamal, A.E., and Schnitzer, M.J. (2013). Long-term dynamics of CA1 hippocampal place codes. Nature Neuroscience 16, 264–266. doi: 10.1038/nn.3329.

## METHODS REFERENCES

Mainali, N., Azeredo da Silveira, R., and Burak, Y. (2025). Universal statistics of hippocampal place fields across species and dimensionalities. Neuron 113, 1110–1120.e3. doi: 10.1016/j.neuron.2025.01.017.

## SUPPLEMENTARY REFERENCES

Bai, Z., and Silverstein, J.W. (2010). Spectral Analysis of Large Dimensional Random Matrices 20. Springer. doi: 10.1007/978-1-4419-0661-8.

Girardeau, G., Benchenane, K., Wiener, S.I., Buzsáki, G., and Zugaro, M.B. (2009). Selective suppression of hippocampal ripples impairs spatial memory. Nature Neuroscience 12, 1222–1223. doi: 10.1038/nn.2384.

Jiang, T. (2021). Wigner’s semicircle law for gaussian random matrices. University of Chicago Mathematics REU. August 2021, 21 pp.

Rasch, B., Büchel, C., Gais, S., and Born, J. (2007). Odor Cues During Slow-Wave Sleep Prompt Declarative Memory Consolidation. Science 315, 1426–1429. doi: 10.1126/science.1138581.

